# Integration of the DD-genome reshapes gene transcription, chromatin architecture and metabolome of allohexaploid wheat leading to enhanced adaptability

**DOI:** 10.1101/2024.10.15.618349

**Authors:** Yanyan Liu, Tao Zhu, Xinkai Zhou, Wei Chen, Chao He, Xin Wang, Chuanye Chen, Jiaqi Wei, Caixia Lan, Mengmeng Liu, Handong Su, Qiang Li, Xin Hu, Siteng Bi, Weizhi Ouyang, Xingwang Li, Hailiang Mao, Masahiro Kishi, Kerstin Kaufmann, Alisdair R. Fernie, Dijun Chen, Wenhao Yan

## Abstract

The integration, through hybridization, of the DD genome into domesticated tetraploid wheat gave rise to allohexaploid wheat, the most cultivated wheat globally growing across diverse environmental conditions. However, the regulatory basis of this integration on increased environmental adaptability in allohexaploid remains largely unexplored. Here, we investigated the change of transcriptome, epigenome as well as the chromatin interactome, and metabolome in three independent polyploidization events representing DD genome integration. Our findings reveal that polyploidization events induce the activation of defense-related genes through comprehensive reorganization of epigenome and chromatin architecture. DD integration not only brings an additional gene copy but also activates the homoeologs existing in the A and B subgenomes through chromatin interactions. Furthermore, secondary metabolites represented by alkaloids and flavonoids that are crucial for environmental adaptation, are significantly enriched following polyploidization. Thus, hexaploid wheat exhibits enhanced tolerance to alkalinity, UV-B light stress and high salt conditions was observed. These results highlight the indispensable role of DD genome integration in the adaptability of allohexaploid wheat during its evolution.

## Introduction

Polyploidy results from whole-genome duplication (WGD) or interspecific hybridization is a widespread genomic phenomenon in eukaryotes (Chen 2007), notably prominent in flowering plants (Leitch and Leitch 2008; Van de Peer et al. 2017). This process leads to an increased genome size and genetic diversity, endowing polyploids with enhanced adaptive plasticity to fluctuating or extreme environments compared with their diploid counterparts. Consequently, polyploidization laid the genetic groundwork for the successful domestication of numerous crops (Leitch and Leitch 2008; Fang and Morrell 2016; Salman-Minkov et al. 2016). Polyploidization is, moreover, crucial for accelerating plant evolution, speciation and biodiversity (Ramsey and Schemske 1998; Otto 2007; Van de Peer et al. 2017). It is proposed that the response to environmental stress plays a pivotal, if not determinative, role in the prosperity of polyploidy (Van de Peer et al. 2021).

Bread wheat (*Triticum aestivum* ssp. *aestivum*, genomes AABBDD) is an allohexaploid with the largest global cultivation area among all crops. Its cultivation area expanded from the Fertile Crescent to almost the terrestrial areas of the globe, encompassing diverse climatic and environmental conditions within just a few millennia following polyploidization (Dubcovsky and Dvorak 2007; International-Wheat-Genome-Sequencing-Consortium 2018; Zhou et al. 2020). The evolutionary journey of bread wheat involved two successive natural hybridization events (International-Wheat-Genome-Sequencing-Consortium 2014). Initially, *T. urartu* (AA genome) intercrossed with a grass believed to be related to *Aegilops speltoides* (SS genome), resulting in the allotetraploid *T. turgidum* (genomes AABB) following genome doubling (Petersen et al. 2006; Avni et al. 2022; Li et al. 2022b). Subsequently, another occasional inter-species hybridization amalgamated the DD genome from diploid *Ae. tauschii* with the genome of *T. turgidum*, giving rise to the ancestral allohexaploid wheat, *T. aestivum* (genomes AABBDD) (International-Wheat-Genome-Sequencing-Consortium 2014), (Wang et al. 2013; Gaurav et al. 2021), which was ultimately domesticated to form bread wheat. There are three distinct lineages of *Ae. tauschii*, lineage1 (L1), lineage2 (L2) and lineage3 (L3), with L2 known as the main origin of wheat D subgenome and L3 is considered to be the other predominant donor of D subgenome in addition to L2. L1 has very limited impact on bread wheat during evolution (Wang et al. 2013; Gaurav et al. 2021; Cavalet-Giorsa et al. 2024).

Allohexaploid wheat shows enhanced tolerance to salt stress, zinc or nitrogen deficiency compared to tetraploid or diploid wheat (Cakmak et al. 1999; Yang et al. 2014; Yang et al. 2018). Recent analyses have revealed that the introgression of genes from wild relatives of wheat played a pivotal role in improved environmental adaptation and agronomic traits during domestication (Jordan et al. 2015; He et al. 2019; Pont et al. 2019; Walkowiak et al. 2020; Zhou et al. 2020; Gaurav et al. 2021). Its response to other stresses, such as alkalinity and excess UV-B radiation, remain unexplored. Furthermore, how the altered genome and gene products such as enzymes or even metabolites contribute to broad adaptability following allopolyploidization is largely unknown. That said, directional genome sequence elimination, rapid expression alteration and dynamic DNA methylation changes during wheat allohexaploidization were found to be involved in the wheat evolution by using either extracted tetraploid wheat or newly synthesized allohexaploid wheat (Shaked et al. 2001; Qi et al. 2012; Li et al. 2014; Zhang et al. 2014; Yuan et al. 2020; Vasudevan et al. 2023). Moreover, a direct link between changes in gene transcription regulatory networks shaped by chromatin status, gene products represented by metabolites and enhanced adaptability has yet to be established.

Changes in epigenetic status are significantly associated with the expression of homoeologous genes in polyploids (Song and Chen 2015). Recent analyses employing techniques such as Hi-C, Hi-ChIP or OCEAN-C approaches to profile histone modifications, open chromatin and three- dimensional (3D) genome structure have underscored the crucial role of regulatory elements and chromatin signatures in governing gene expression and the asymmetric gene transcription in wheat (Li et al. 2019b; Concia et al. 2020; Jia et al. 2021; Wang et al. 2021; Pei et al. 2022; Yuan et al. 2022; Zhao et al. 2023). However, the level of reorganization of both epigenetic landscape and three- dimensional chromatin interactions upon the introduction of the DD genome during wheat evolution remains unclear. Moreover, how such changes subsequently influenced gene transcription to produce variable content of metabolites is also currently unknown. Our understanding of polyploidization has primarily focused on genomic variation or gene transcription dynamics and the knowledge of downstream features such as metabolites is lacking. Metabolites are both pivotal determinants of crop quality traits and actors in stress responses. For instance, flavonoids are known to play a critical role in abiotic stress responses (Yonekura-Sakakibara et al. 2014; Peng et al. 2017; Chen et al. 2018; Fang et al. 2019), while terpenoids contribute to environmental adaptability and fruit quality (Shang et al. 2014; Huang et al. 2019).

Synthetic hexaploid wheat (SHW) is generated through the crossing of tetraploid wheat *T. turgidum* with its diploid relatives *Ae. tauschii*, followed by artificial chromosome doubling of the triploid F_1_ plants (Li et al. 2018a; Rosyara et al. 2019). As such generation of SHW simulates the hexaploidization process (Wan et al. 2022). Here, a cohort of SHWs and their parental tetraploid wheat and diploid *Ae. tauschii* were used to systematically investigate how the DD-integration mediated hexaploidization events enhance wheat adaptability by altering the transcriptome, epigenome, 3D chromatin interactome and metabolome. We focused on the activation of genes and metabolic pathways associated with environmental adaptability paying particular attention to resistance to alkaline, salt and UV-B stress during polyploidization.

## Results

### The experimental system

Hexaploid wheat, is the most extensively cultivated crop across diverse global regions characterized by diverse environmental conditions (**Supplementary Fig. S1**), indicating its remarkable adaptation and versatility. To elucidate the potential impact of DD integration within environmental adaptability, we conducted a comprehensive multi-omics analysis using a set of genotypes comprising three synthetic hexaploid wheats (SHW3, SHW4 and SHW5), their common tetraploid parent (*T. turgidum* ssp. *Durum*, AABB#7) and their independent diploid parents (*Ae. tauschii*, DD#10, DD#11 and DD#12) (**Fig. 1A**; **Supplementary Data Set 1**). Notably, these diploid parents belong to two distinct DD lineages, with DD#10 and DD#11 originating from lineage 2, which has been introduced into current bread wheat, while DD#12 comes from lineage 1, a lineage hardly found in bread wheat (Wang et al. 2013; Gaurav et al. 2021) (**Supplementary Fig. S2A**). Fluorescent *in situ* hybridization (FISH) and genomic *in situ* hybridization (GISH) analyses unequivocally confirmed that all SHWs used in our study possess 42 chromosomes (2n = 6× = 42), while the tetraploid and diploid parents possess 28 (2n = 4× = 28) and 14 (2n = 14) chromosomes, respectively (**Supplementary Fig. S2, B to H**).

**Figure 1.**
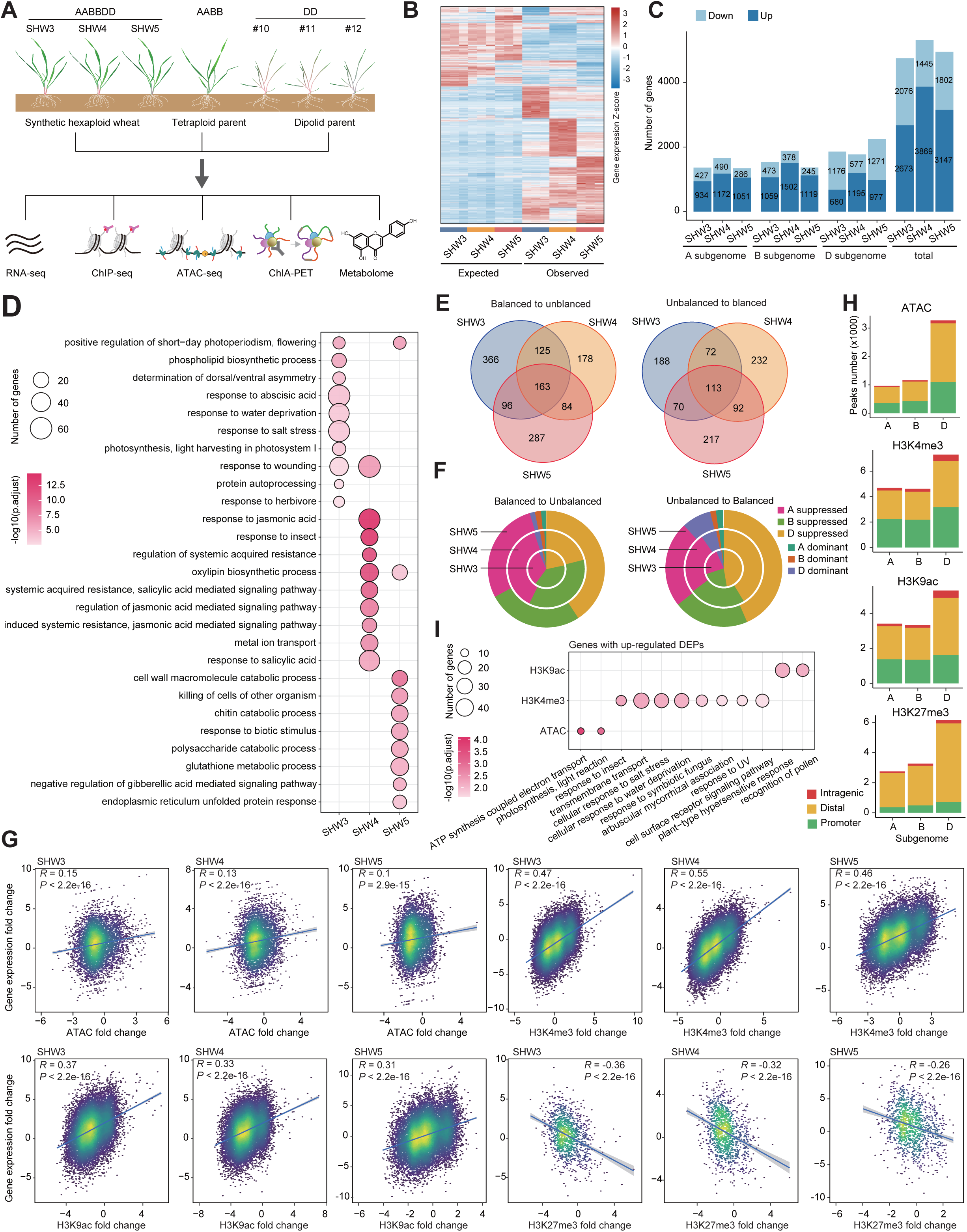
Reshaping of transcriptome during the DD-integration mediated polyploidization. **A)** Schematic outlining the experimental design of multi-omics analysis for DD genome integration mediated polyploidization events. The fully expanded leaves from 21-days wheat seedlings were collected for RNA-seq, ChIP-seq, ATAC-seq, ChIA-PET and metabolome analysis. **B)** Heatmap showing identified differentially expressed genes (DEGs) during wheat polyploidization events. **C)** Number of DEGs affected by the polyploidization in different subgenomes of SHWs. Activated genes and suppressed genes were indicated respectively by dark blue and light blue bars. **D)** Top ten GO terms enriched in up-regulated. Dot size represents the gene number enriched in GO terms and color gradation represents significance of enrichment calculated as –log_10_ (FDR). **E)** Numbers and overlapping of triads that changed asymmetric expression patterns (left, balanced to unbalanced; right, unbalanced to balanced) during the polyploidization events. **F)** Proportion of categories that altered its balance status during polyploidization. **G)** The relationship between gene expression fold change with peak fold change of ATAC-seq and ChIP-seq of H3K4me3, H3K9ac and H3K27me3. **H)** The number of shared differentially enriched peak (DEP) in SHWs for ATAC-seq, ChIP-seq of H3K4me3, H3K9ac and H3K27me3 in different subgenomes. **I)** Bubble plot showing the top enriched GO terms of genes with up-regulated chromatin accessibility and histone modifications of H3K4me3 or H3K9ac during wheat polyploidizations. The size of dots indicates number of genes in the terms, and the gradation of colors represents the significance as –log_10_ (FDR).

### Massive activation of adaptability genes during DD genome integration

We generated high-quality gene transcription profiles for all genotypes (**Supplementary Fig. S3** and **Data Set 2**). The total number of expressed genes (TPM > 0.5) among subgenomes was comparable (**Supplementary Fig. S4A**). In order to delineate changes in gene expression following polyploidization, we conducted a theoretical gene expression profile by aggregating gene expression from the corresponding parental tetraploid and diploid to create an “expected” gene profile from *in-silico* hexaploid wheat (Ramírez-González et al. 2018) . This approach assumes no interactions or coordination among the AB and D subgenomes upon their amalgamation into the hexaploid wheat cell. Meanwhile, the gene expression observed through RNA-seq analysis in SHWs was designated as “observed” gene expression.

We compared the “observed” gene expression in SHWs with the theoretical “expected” expression and identified 9830 differentially expressed genes (DEGs) that are supposed to be influenced by polyploidization in at least one of the SHWs (**Fig. 1B**). Among these, SHW3, SHW4 and SHW5 exhibited 4749, 5314 and 4949 DEGs, respectively, with 2673-3869 up-regulated and 1445-2076 down-regulated genes (**Fig. 1C**; **Supplementary Data Sets 3** to **5**). Furthermore, 245- 490 genes were down-regulated in the A and B subgenomes, whereas this number increased to 577- 1271 in the D subgenome. The number of up-regulated genes was comparable among the A, B and D subgenomes (**Fig. 1C**). A significant overlap of DEGs were observed across different SHWs for both up-regulated and down-regulated genes (**Supplementary Fig. S4, B and C**). The up-regulated DEGs were predominantly related to various aspects of environmental adaptation, including functional terms related to multiple stimulus response across the A, B, D subgenomes of different SHWs (**Fig. 1D**; **Supplementary Fig. S5A**). Interestingly, the “translation” related GO terms were commonly enriched in the down-regulated genes, particularly those from the A and B subgenomes (**Supplementary Fig. S4D** and **S5B**). These results suggest that wheat polyploidization can activate adaptation genes in both the original A and B subgenomes as well as within the newly integrated D genome itself, while simultaneously requiring the re-balancing of activities of genes related to fundamental processes.

We categorized homoeolog triads into seven expression patterns: balanced, A- /B- /D-suppressed or A- /B- /D- dominant (Ramírez-González et al. 2018). The expressed triads (the sum TPM of A, B, D subgenome homoeologs > 0.5) were analysed and we observed high correlation among genes within the same category across the three SHWs (**Supplementary Fig. S6A**). The ratios of homoeolog-dominant and D-suppressed decreased following polyploidization, while the ratios of A- and B-suppressed and balanced category increased in the SHWs compared with its parents (**Supplementary Fig. S6, B to D**). We further categorized homoeologous genes with changed expression between “observed” and “expected” datasets into two types, “balanced-to- unbalanced” and “unbalanced-to-balanced”. The former represents triads that changed from symmetrical (balanced) to asymmetrical (unbalanced, homoeolog-suppressed or homoeolog-dominant) expression patterns after polyploidization (**Supplementary Data Sets 6 to 8**), while the latter indicates triads that shifted from unbalanced to balanced expression in SHWs (**Supplementary Data Sets 9 to 11**). The altered proportions of categories between “expected” and “observed” groups were the result of orchestration within homoeologs (**Supplementary Fig. S7, A to C**). Among the expressed triads in SHWs, there are 550-750 and 443-509 triads showed altered pattern from balanced to unbalanced and from unbalanced to balanced types, respectively (**Fig. 1E**). Interestingly, most of the balanced-to-unbalanced triads tended to be constitutively expressed across various tissue types (**Supplementary Data Set 12**), whereas the unbalanced-to-balanced triads exhibited both tissue-specific and constitutive expression patterns (**Supplementary Fig. S7, D and E**). Furthermore, almost all balanced-to-unbalanced triads were homoeolog-suppressed and only 0.9%-2.9% of triads becoming homoeolog-dominant in SHWs (**Fig. 1F**; **Supplementary Fig. S8, A to C** and **Data Sets 6 to 8**). Conversely, for the unbalanced-to-balanced type, a significant proportion of unbalanced categories that became balanced after DD integration were homoeolog-suppressed, while only 0.9% to 2.0% of triads for A- or B- dominant and 4.4% to 8.1% for D-dominant patterns changed to balanced state following polyploidization (**Fig. 1F**; **Supplementary Fig. S8, D to F** and **Data Sets 9 to 11**). These results collectively demonstrated the coordination of gene transcription among A, B and D subgenomes after polyploidization and suggested the balance of homoeolog gene expression levels following the integration of the DD genome, particularly through the reduction of asymmetric expression for genes in the DD subgenome.

### Altered histone modifications and chromatin accessibility during wheat allohexaploidization correlate with transcriptional changes of genes linked to environmental adaptability

Gene transcription is highly related with chromatin status, including histone modifications and chromatin accessibility. To study how the integration of the DD genome reshapes the epigenome to affect gene transcription, we mapped genome-wide histone modifications of H3K4me3, H3K9ac, H3K27me3 and estimated the chromatin accessibility in the SHWs and their tetraploid and diploid parents (**Supplementary Data Set 13**). Correlation analysis revealed a significant positive relationship between changes in gene expression and chromatin accessibility (ATAC-seq) and H3K4me3 or H3K9ac modifications, while a negative correlation was observed with H3K27me3 (**Fig. 1G**).

We next compared the “expected” and “observed” histone modification and chromatin accessibility status following polyploidization and observed a significant decrease in chromatin accessibility in the D subgenome after polyploidization in all three SHWs (**Supplementary Fig. S9**). The H3K27me3 level was higher after polyploidization in all subgenomes of the three SHWs, while the levels of H3K4me3 and H3K9ac remained comparable before and after polyploidization (**Supplementary Fig. S9**). Additionally, we identified differentially enriched peaks (DEPs) derived from ATAC-seq (corresponding to accessible genomic regions) and ChIP-seq following polyploidization (**Supplementary Data Sets 14 to 17**). Remarkably, more DEPs were identified in the D subgenome than in the A or B subgenomes (**Fig. 1H**). In addition, GO analysis using genes with DEPs in their promoters or intragenic regions revealed that genes with up-regulated DEPs were enriched with terms related to environmental/external stimulus response, such as “cellular response to salt stress”, “response to UV” and “plant-type hypersensitive response” (**Fig. 1I**), while genes with down-regulated DEPs were predominantly associated with “photosynthesis” or “translation” (**Supplementary Fig. S10**). These results underscore the significance of changes of histone modification and chromatin accessibility in mediating coordinated expression between the D subgenome and the original A and B subgenomes. In particular, dynamic changes of chromatin status appears to be one of the major regulatory mechanisms underlying the altered expression of those genes related to environmental adaptability.

### Reorganization of 3D chromatin architecture may serve as the underlying driving force to reshape gene transcription during DD genome integration

Collective evidence suggests that long-range chromatin interactions have significant impact on gene transcription (Tang et al. 2015; Li et al. 2019a; Peng et al. 2019; Zhao et al. 2019; Deng et al. 2022). Given the close association of H3K4me3 with gene transcription and the enrichment of H3K4me3- modified genes with various GO terms (**Fig. 1, G and I**; **Supplementary Fig. S10**), H3K4me3 modification can be used as an indicative marker of gene transcription in order to explore the impact of DD genome integration-mediated allopolyploidization in wheat. To examine the changes of H3K4me3-associated chromatin interaction during wheat polyploidization, we performed long-read ChIA-PET associated with H3K4me3 in SHW3 and its tetraploid and diploid parents (**Fig. 2, A to C**; **Supplementary Data Set 18**). The highest number of H3K4me3-associated chromatin interactions were detected within the D subgenome, followed by the A subgenome, with the least interactions occurring within the B subgenome in SHW3 (**Fig. 2A**; **Supplementary Fig. S11A**). This observation is consistent with published Hi-C data (Concia et al. 2020; Jia et al. 2021). Chromatin interactions within the A subgenome of the tetraploid occur at higher frequency than within the B subgenome (**Fig. 2B**; **Supplementary Fig. S11B**), indicating that the difference between the A and B subgenome in hexaploid wheat (SHW3) were rooted in their tetraploid progenitor. Besides, we detected more loops among homeologues than non-homeologues, with the highest interaction frequency observed in chromosome group 2 (2A, 2B and 2D) and group 5 but the lowest in chromosome group 4 (**Supplementary Fig. S11, C and D**).

**Figure 2.**
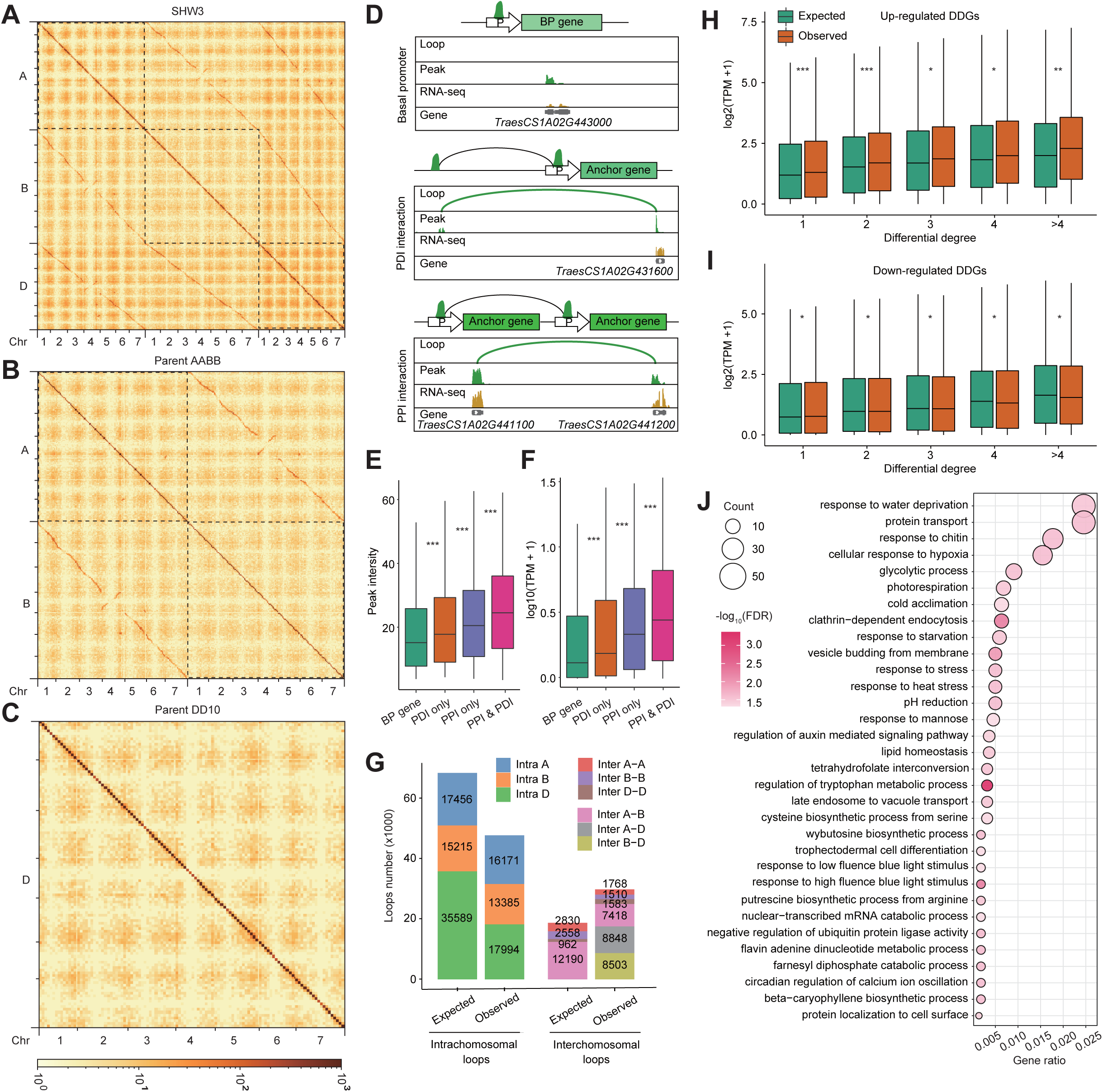
Dynamics of H3K4me3-associated chromatin architecture following allohexaploidization event. **A-C)** ChIA-PET heatmap showing H3K4me3-associated chromatin interactions in SHW3 **A)** and its tetraploid **B)**, diploid **C)** parents. **D)** Schematic models of H3K4me3-associated chromatin interactions. BP model, genes with H3K4me3 modification but are not involved in chromatin interactions. PPI model, anchor genes involved in H3K4me3-associated proximal interactions. PDI, anchor genes involved in interactions between H3K4me3-associated proximal and distal modification region. **E, F)** Boxplot showing the intensity of H3K4me3 modification in different chromatin models **E)** and the expression level of BP genes and genes involved in PDI, PPI interactions **F)** in SHW3. The gene expression levels are shown as log_10_(tpm + 1). \*\*\**P* < 0.001. **G)** Comparation of the intrachromosomal and interchromosomal loops in SHW3 before and after polyploidization. **H, I)** Boxplots showing the expression level of up-regulated DDGs **H)** and down-regulated DDGs **I)** in expected and observed dataset of SHW3. \**P* < 0.05, \*\**P* < 0.01, \*\*\**P* < 0.001. **J)** Bubble diagram showing enriched GO terms of up-regulated DDGs in SHW3. The dot size represents the number of genes in the corresponding GO term, and the gradation of color indicates the significance of terms as –log_10_ (FDR).

We refer to genes with chromatin loops in their promoter as anchor genes, and genes with H3K4me3 modification but without chromatin loops being defined as BP genes (genes with a basal promoter) (**Fig. 2D**). Depending on whether the chromatin-loop involved peaks were located in the proximal (3.5 kb upstream of transcription start site [TSS] to transcription end site [TES]) or distal genic region (genomic regions out of proximal region), chromatin interactions were divided into proximal-proximal interactions (PPIs) and proximal-distal interactions (PDIs) (**Fig. 2D**). Compared with BP genes, anchor genes exhibited significantly higher H3K4me3 levels in their promoters and showed higher expression levels. Indeed, anchor genes with both PPI and PDI chromatin interactions displayed the highest H3K4me3 modification intensity and expression levels among all wheat genes (**Fig. 2, E and F**). Moreover, anchor genes with a higher degree of interaction (more connected loops), possessed a higher median H3K4me3 peak intensity in their promoter and displayed a higher expression level (**Supplementary Fig. S12, A and B**), indicating a positive correlation of chromatin interaction degree with both H3K4me3 modification and gene expression levels. The above results collectively suggest a regulatory role of chromatin interactions in modulating gene transcriptional activities in wheat.

We next generated a virtual “expected” chromatin interaction map based on the chromatin interactions in tetraploid and diploid parents and then compared it with that in SHW3 (designated as “observed” dataset) to estimate the impact of altered chromatin interactions on gene transcription during DD integration. We found that in the D subgenome, the observed genomic span of intrachromosomal loops were shorter than the expected loops span following polyploidization, but no obvious difference was observed in the A and B subgenomes (**Supplementary Fig. S12, C to E**), suggesting a more intensive reorganization of the DD genome compared with the AA and BB genomes after DD integration. In total, we observed 77180 interaction loops connecting 46445 anchor genes in SHW3 in comparison with 86800 interactions and 50605 anchor genes identified in the “expected” dataset (**Supplementary Fig. S12, F and G** and **Data Set 18**). Notably, 47901 loops were exclusively found before allopolyploidization, while 38281 loops were specifically detected following the DD genome integration (**Supplementary Fig. S12F** and **Data Sets 19 to 20**). It is remarkable that more interchromosomal interactions were produced following the DD-genome integration. This is likely due to the combinatorial effects of the reduction of intrachromosomal interactions in the DD subgenome and the interchromosomal interactions between A and B subgenomes and the increase of newly established interactions between A-D and B-D subgenome (**Fig. 2G**).

Next, we compared the “observed” interaction degrees of all anchor genes with those “expected” in order to identify differential degree genes (DDGs). A total of 17356 genes exhibited increased degrees (up-regulated DDGs), including 3892 genes that changed from BP genes to anchor genes. Conversely, 24902 genes displayed decreased degree of interactions (down-regulated DDGs), including 8052 genes that shifted from anchor genes to BP genes following polyploidization (**Supplementary Figs. 12G, 13A, B** and **Data Set 21**). While up-regulated DDGs were distributed equally across the A and B subgenomes but were less abundant in the D subgenome, more down- regulated DDGs were found in the D subgenome than in the A and B subgenomes (**Supplementary Fig. S13, A and B**). These up-regulated DDGs showed a significantly higher expression level, whereas down-regulated DDGs showed a significantly lower expression level in the “observed” dataset (**Fig. 2, H and J**). We next conducted GO analysis using DDGs with differential interaction degree ≥ 3, and a total of 3397 up-regulated DDGs and 4971 down-regulated DDGs were detected (**Supplementary Data Set 21**). In addition to the genes for basic physiological processes such as protein transport and glycolytic process, genes for response to environmental stimuli such as water deprivation, chitin, cold acclimation, PH reduction and heat stress are significantly enriched in up- regulated DDGs but not in down-regulated DDGs (**Fig. 2J**; **Supplementary Fig. S13C** and **Data Set 22**).

### Activation of adaptive genes through enhanced chromatin interaction facilitates environmental adaptability following allopolyploidization

To visually represent the chromatin architecture affected by DD genome integration, chromatin interaction networks were constructed using 1545 core DEGs that showed altered H3K4m3 modification or with differential interaction degree ≥ 3. Interestingly, most of these genes were not connected with each other in the parental AABB or DD genomes (**Fig. 3, A and B**), but after polyploidization, they were tightly linked to form a large regulatory module following DD genome integration (**Fig. 3C**). For example, a group of receptor-like kinases (RLKs) encoded genes with roles in sensing diverse environmental changes and involvement in biological process related to biotic and abiotic stress (Morris and Walker 2003; Tang et al. 2017; Shumayla et al. 2019), displayed increased interaction degrees, enhanced H3K4me3 modification, and elevated expression following DD genome integration (**Fig. 3, D and F**). Furthermore, a wheat chitinase encoded gene, *TaCHI-2B* (*TraesCS2B02G047300*), which is involved in the plant defense response by hydrolyzing chitin (Grover 2012; Vaghela et al. 2022), formed chromatin loops with other eight genomic regions in the “observed” dataset compared with only one loop in the “expected” data and was accompanied by increases level of histone modification and transcriptional activation following polyploidization (**Fig. 3, E and F**). These results collectively underscore the alteration of gene expression via 3D chromatin interaction following the wheat DD integration event, serving as a significant driving force to shape gene transcription among subgenomes in polyploidy species.

**Figure 3.**
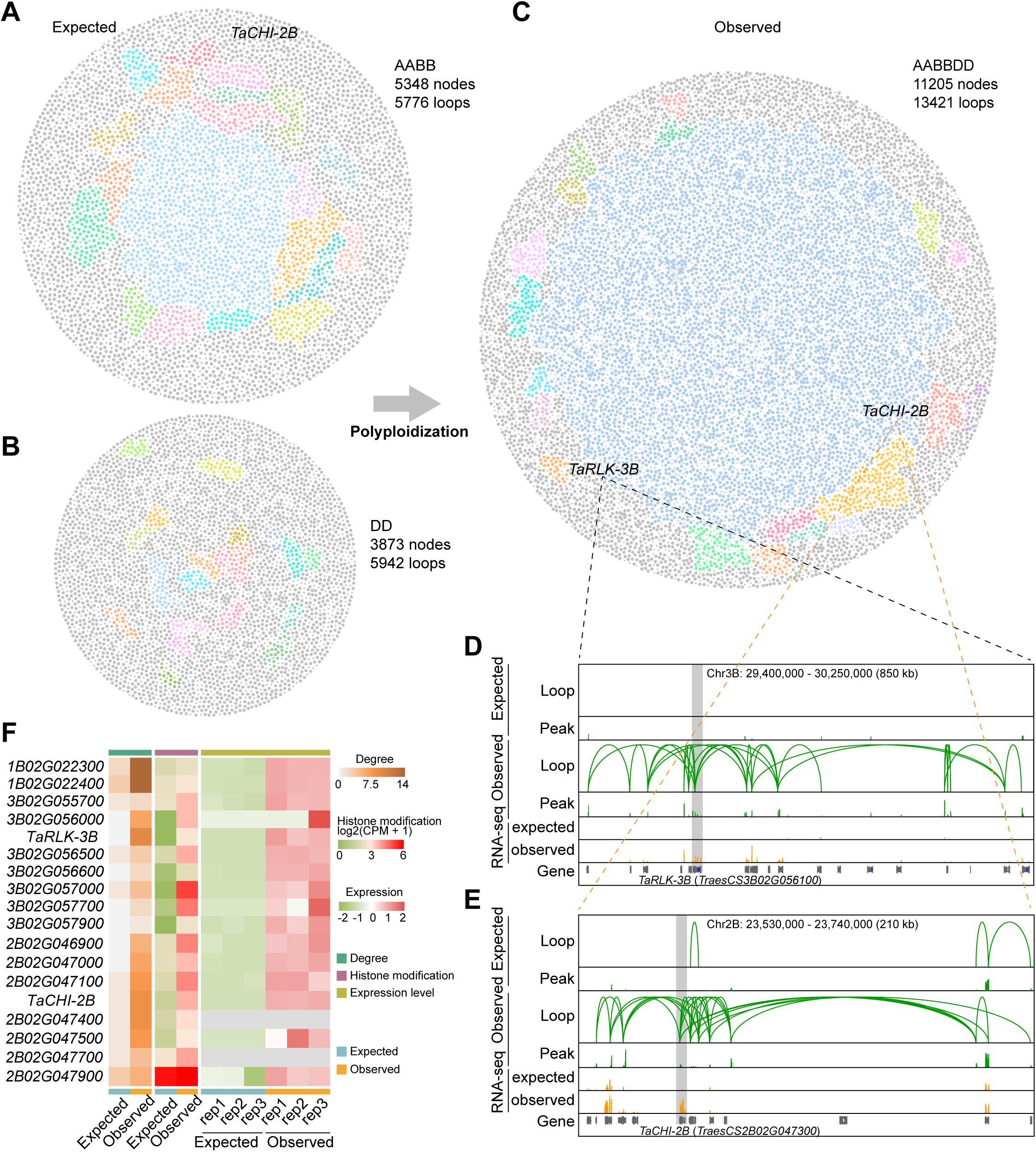
Dynamics of H3K4me3-associated chromatin architecture following wheat allohexaploidization event. **A-C)** Chromatin interaction networks of the core 1545 DEGs in SHW3 **C)** and its tetraploid **A)** and diploid **B)** parents. The top 20 modules are presented using different color. **D)** A browser screenshot showing the H3K4me3-associated chromatin interactions, modification peaks and RNA-seq data in an active region containing tandem RLK-encoding genes before and after polyploidization. **E)** A browser screenshot showing comparation of chromatin interactions, ChIP-seq peaks and RNA-seq data in the actively transcribed region of *TaCHI-2B* (*TraesCS2B02G047300*) before and after polyploidization. **F)** Heatmap showing increased chromatin interaction degree, enhanced histone modification, and activated transcription of the genes following DD-integration.

To elucidate potential mechanisms underlying the enhancement of adaptability through polyploidization, we collected adaptative genes from published studies and identified wheat homologs of *Arabidopsis* genes (**Supplementary Data Set 23**) before constructing a gene network by considering chromatin interaction. Our analysis revealed numerous genes exhibiting increased chromatin spatial interaction frequency and elevated expression levels following polyploidization, including genes related to UV-B tolerance, alkaline tolerance, salt tolerance, cold resistance, nitrogen assimilation, as well as the flavonoid biosynthesis pathway (**Fig. 4, A to C**). For example, *TaCCD1* and *TaHA2*, known as positive regulators in wheat alkali stress tolerance (Cui et al. 2023), showed enhanced interaction degree in “observed” networks accompanied by increased transcription following DD genome integration. Similarly, *TaHY5-like*, *TaTDP1*, and *TaERF4*, wheat homologs of *Arabidopsis HY5*, *TDP1*, and *ERF4* identified in Tibetan semi-wild wheat for their adaptation to UV-B radiation and cold acclimation in high-altitude environments (Guo et al. 2020), exhibited obviously increased chromatin interaction frequency in the allohexaploid wheat. Conversely, *TaPP2C.D1* and *TaPP2C.D2* (Cui et al. 2023), negative regulators of alkali resistance, demonstrated decreased chromatin interaction following polyploidization (**Fig. 4, A to C**).

**Figure 4.**
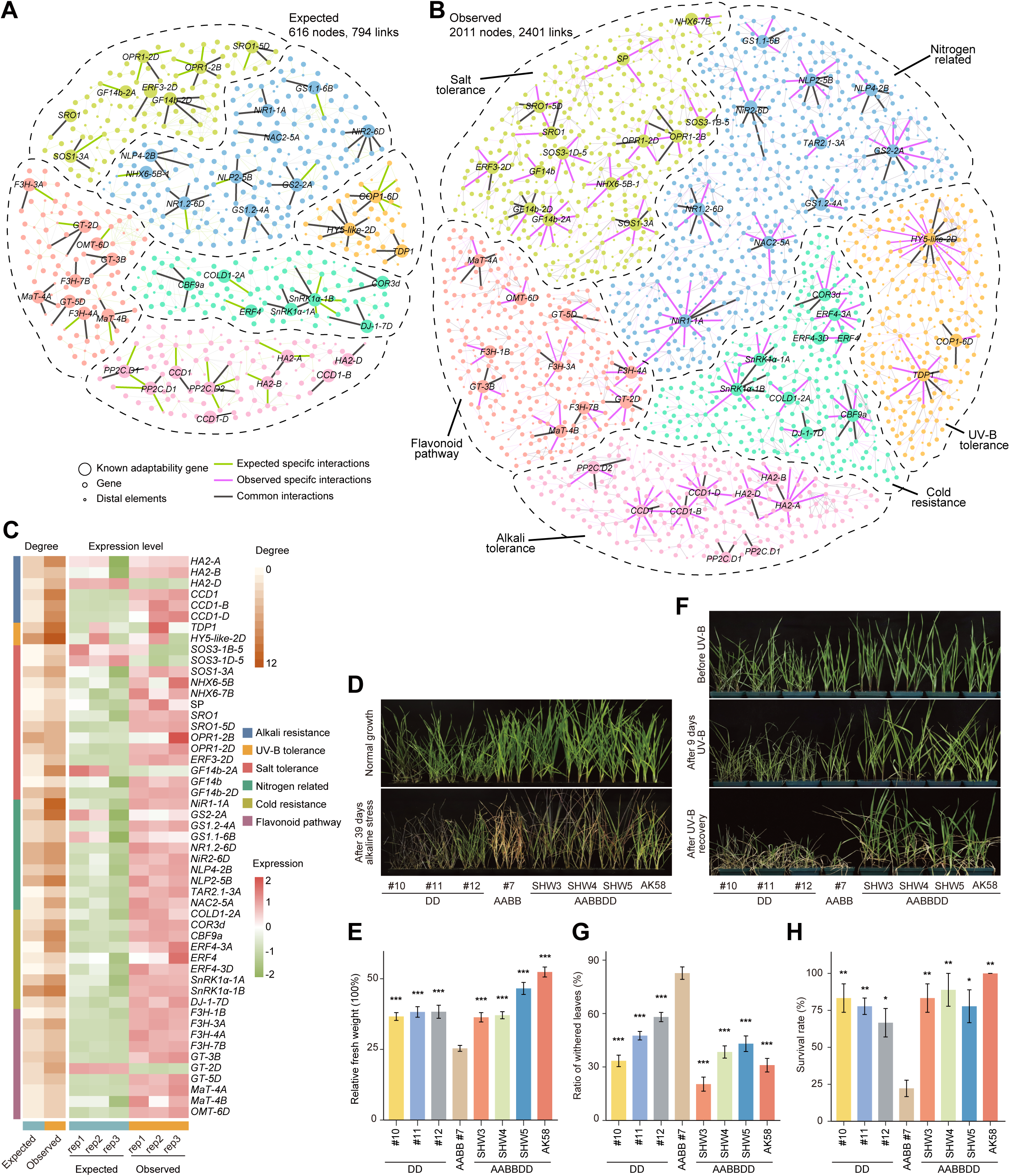
Activation of adaptability genes through enhanced H3K4me3-associated chromatin interactions after polyploidization. **A, B)** chromatin interaction networks built using adaptability genes before **A)** and after **B)** polyploidization. Different dot colors represent nodes from different networks of UV-B tolerance, alkali tolerance, salt tolerance, cold resistance, nitrogen related genes, and flavonoid pathway. The light green lines represent specific interactions before polyploidization. The light purple lines indicate newly appeared interactions after DD-genome integration. The black lines indicate common interactions both in before and after networks. **C)** Heatmap showing increased interaction frequency (degree) and enhanced expression level of the adaptation related genes. **D-H)** Phenotypes of AK58, SHWs, tetraploid and diploid parents (AABB and DD) under alkaline stress **D, E)** and UV-B treatment **F-H)**. \**P* < 0.05, \*\**P* < 0.01, \*\*\**P* < 0.001.

Previous studies have showed the enhanced fitness of allohexaploid wheat to salt stress and nitrogen or zinc deficiency conditions compared with its tetraploid or diploid progenitors (Cakmak et al. 1999; Yang et al. 2014; Yang et al. 2018). To further investigate the potential adaptability of allohexaploid wheat to additional stress conditions, we applied stress to the three SHWs (SHW3, SHW4 and SHW5), their shared tetraploid parents and independent diploid parents, with the widely grown allohexaploid wheat variety Aikang 58 (AK58) (Jia et al. 2023) serving as a control. When all plants were irrigated with 250 mM NaCl solution, allohexaploid wheat performed better than its tetraploid and diploid parents, with 11.7% to 15.5% relative fresh weight of salt-treated hexaploid plants comparing with 9.5% in tetraploidy and 1.5% to 5.3% in diploidy plants after 43 days treatment (**Supplementary Fig. S14, A and B**), consistent with previous research (Yang et al. 2014). Under 100 mM mixed alkaline stress, tetraploid wheat showed obvious growth inhibition with more whitened/withered leaves than observed in the hexaploid (including the three SHWs and AK58) and diploid after 39 days of irrigation by water containing 100 mM alkaline (**Fig. 4D**). The relative fresh weight of alkaline-treated tetraploid plants is only 25.4% of the corresponding plants growth in normal conditions, significantly lower than alkaline-treated hexaploid and diploid wheat which exhibited relative fresh weights of 36.4% to 52.4% and 36.6% to 38.3% respectively (**Fig. 4E**). When subjected to UV-B stress, 82.7% leaves of tetraploid wheat withered after 9 days and 22.2% tetraploid plants survived after returning to normal condition for 14 days. By contrast, allohexaploid and diploid plants showed a lower proportion of withered leaves (20.4% to 43.1% in allohexaploid and 33.4% to 58.1% in diploid) after UV-B exposure and 77.8% to 100% and 66.7% to 83.3% of the allohexaploid and diploid plants survived respectively after the cessation of the stress (**Fig. 4, G and H**). These results showed the clearly stronger tolerance and growth advantage of hexaploid wheat under alkaline, UV-B, and salt stress conditions than their tetraploid progenitors. They were furthermore in keeping with the observed transcription of genes related with adaptability mentioned above.

### DD-integration mediated polyploidization led to comprehensive alterations in metabolome

Our results from gene transcription and gene regulatory network analysis revealed that one of the most significant features is the enhanced expression of genes related to environmental adaptability (**Fig. 1, D and I; Fig. 2J** and **4**). Metabolites, especially secondary metabolites, are crucial in plant responses to environmental stimuli, they thereby influence adaptability (Huang et al. 2019; Yu et al. 2020). Domestication has been demonstrated to profoundly alter metabolite levels in various species, including tetraploid wheat, potato and radish (Beleggia et al. 2016; Fasano et al. 2016; Zhang et al. 2021b). To explore metabolome changes following the DD genome integration, we conducted liquid chromatography-tandem mass spectrometry (LC-MS/MS) analysis on the allohexaploid SHWs and their tetraploid and diploid parents (**Supplementary Data Set 1**) (Chen et al. 2013). A total of 839 metabolites were detected and 477 metabolites were putatively annotated, including 84 alkaloids, 76 amino acids and peptides, 64 carbohydrates, 84 fatty acids, 151 phenylpropanoids and 18 terpenoids (**Supplementary Data Set 24**).

Principal component analysis (PCA) using these 839 detected metabolites clearly distinguished the allohexaploid SHWs from their tetraploid and diploid parents. Notable differences were observed in DD#12 from DD#10 and DD#11, and SHW5 from SHW3 and SHW4, respectively (**Fig. 5A**), in line with the fact that DD#12 belongs to lineage 1 (**Supplementary Fig. S2A**). Metabolites were further categorized into eight clusters based on their relative contents across different genotypes. Allohexaploid SHWs exhibited the highest content of metabolites in cluster 1 and cluster 2, whereas cluster 3 had the highest content in tetraploid wheat, and clusters 4-6 had higher content in *Ae. tauschii* (genomes DD) than any polyploids (**Fig. 5B**). Alkaloids were more prevalent in cluster 1 and 2, while carbohydrates were prominent in cluster 3, and fatty acids were enriched in clusters 4-6 (**Fig. 5C**). All the above results indicate extensive changes in metabolite levels during polyploidization.

**Figure 5.**
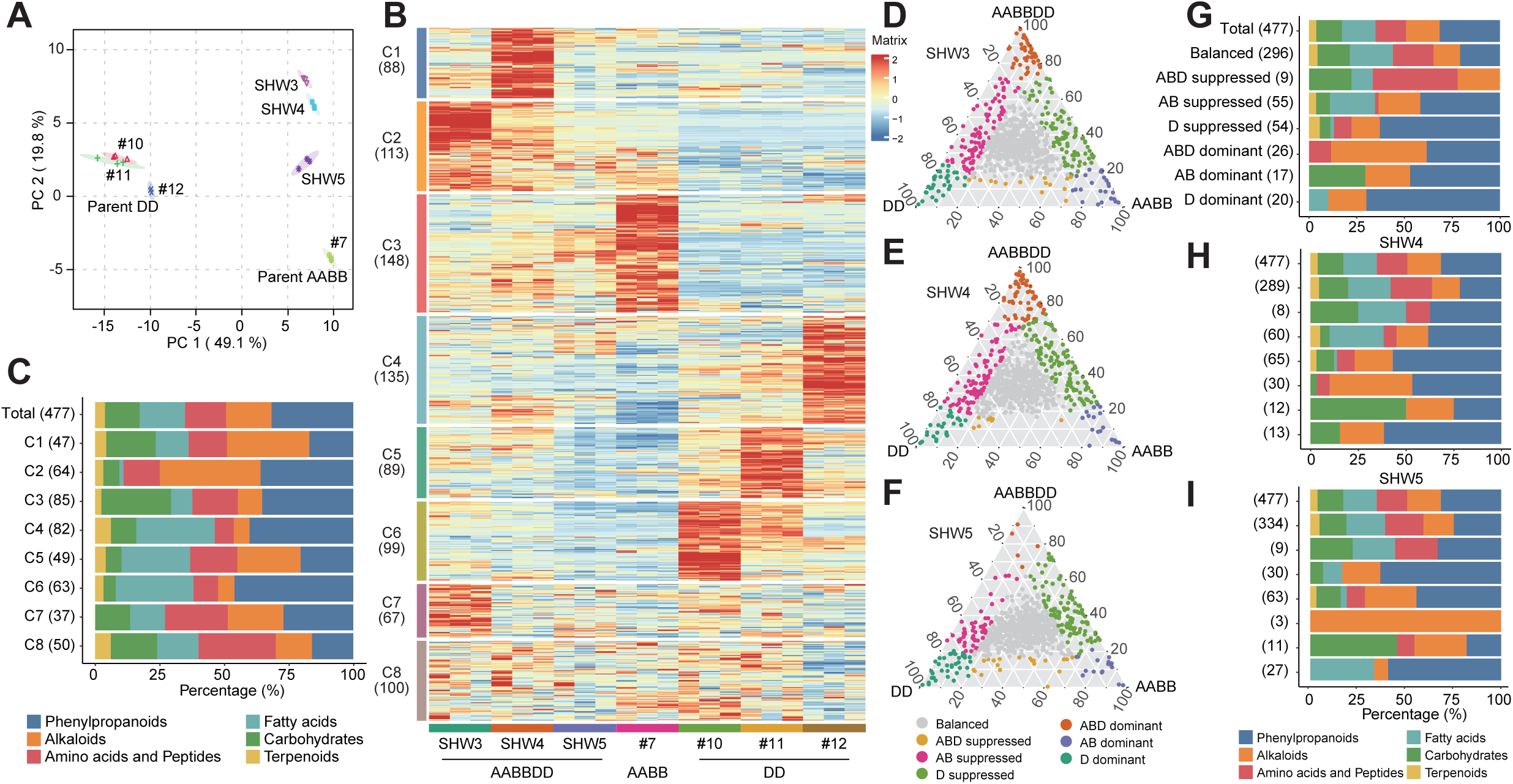
Metabolome analysis of allohexaploid SHWs and the tetraploid, diploid parents. **A)** Principal component analysis (PCA) of metabolome data generated from SHWs and its parents. Different colors and shapes represent different samples, *n* = 3. **B)** Heatmap showing eight major clusters grouped based on the contents of a total of 839 metabolites in allohexaploid SHWs, tetraploid and diploid parents. The number of metabolites in each cluster is given in parentheses. **C)** Proportions of metabolite classes with annotated structures within the eight clusters. Different colors indicate the six different metabolite classes. The proportions of all the 472 metabolites in the six classes with known structures is shown. **D-F)** Ternary plot showing diversity of metabolites between SHW (AABBDD) and its tetraploid (AABB), diploid (DD) parents of SHW3 **D)**, SHW4 **E)**, SHW5 **F)**. The balanced type of metabolites that showed similar content in the three materials are shown as grey dots, and the unbalanced type are shown as colorful dots. **G-I)** Proportions of metabolite classes with annotated structures within the seven categories shown in **D-F)** of SHW3 **G)**, SHW4 **H)**, SHW5 **I)**.

Metabolites were classified into seven categories analogous to the asymmetric gene expression analysis to study their dynamic changes during polyploidization (**Fig. 5, D to F**). Approximately 64.8% to 71.3% of metabolites exhibited a balanced pattern across SHWs and their parents. Notably, alkaloids and phenylpropanoids showed the most significant changes (**Supplementary Fig. S15, A to C**). In addition, alkaloids are enriched in the ABD dominant categories in SHWs and carbohydrates is prominent in AB dominant categories (**Fig. 5, G to I**). These results suggest that DD genome integration-mediated polyploidization led to reduced levels of carbohydrates and increased levels of secondary metabolites, particularly alkaloids, in allohexaploid SHWs.

### Activation of flavonoid and benzoxazinoid pathways during DD genome integration

Flavonoids are known for their roles in response to various stresses such as oxidative and drought stress, UV light and pathogen infections (Yonekura-Sakakibara et al. 2014; Peng et al. 2017; Chen et al. 2018; Fang et al. 2019). In plants, flavonoids are usually stored as glycosides in vacuoles, with the activities of glycosyl transferases being a key step in their accumulation (Zhao and Dixon 2010; Ren et al. 2020). We noticed that several glycosyl derivatives of flavonoid synthesis were present at very low levels in AABB genomes but at high level in both DD and AABBDD genomes (**Fig. 6, A and B**), suggesting that the increased content of flavonoids in SHWs was resulted from the DD genome integration. Strikingly, the expression levels of the DD genome orthologs of rice flavonoid biosynthesis genes were significantly higher than those of A and B members both before and after the DD genome integration (**Fig. 6C**). This higher expression level was attributed to the synergistic effect between active and negative epigenetic markers (**Supplementary Fig. S16, A to C**).

**Figure 6.**
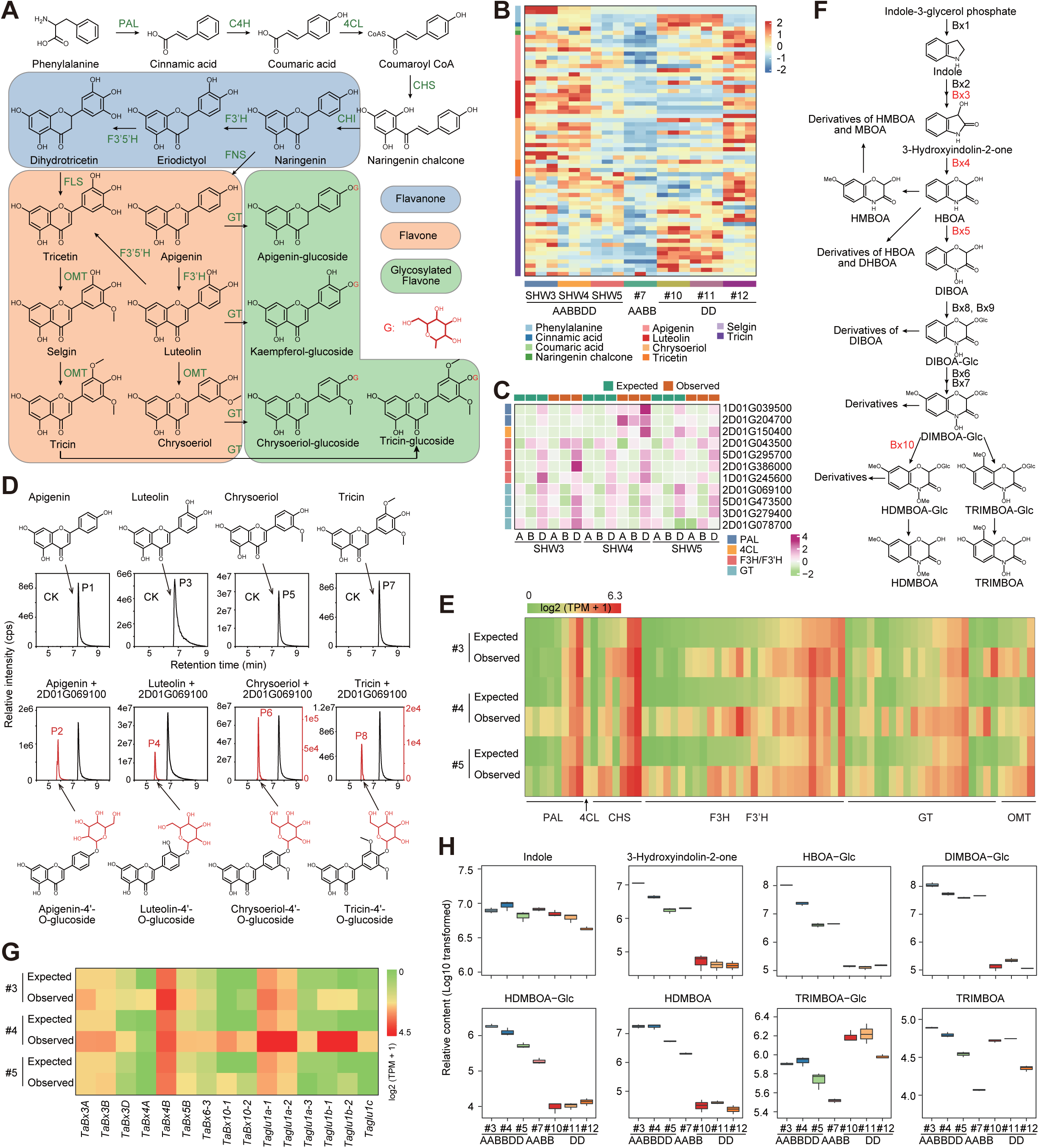
Integration of excellent genes from DD genome optimized flavonoid and benzoxazinoid pathways of allohexaploid wheat. **A)** The core flavonoid pathway in plants. The different colored shadow indicates different classes of flavonoids. **B)** Heatmap showing flavonoid pathway metabolites with high contents in allohexaploid SHWs (AABBDD) compared with its tetraploid parent (AABB). Different colors on the left of the heatmap indicate core flavonoid metabolites. **C)** Putative flavonoid pathway triad genes whose D homoeolog showed highest expression level compared with A and B homoeologs. **D)** Biochemical function of O-glucosyltransferase TraesCS2D01G069100 on flavonoids. The red peaks indicate newly produced metabolites in the reactions; glycosyls newly transferred to flavonoid substrate were represented by red. **E)** Heatmap showing putative flavonoid pathway genes with activated expression after DD-integration. Expression level is given as log_2_(tpm + 1). **F)** Biosynthesis pathway of benzoxazinoids (BXs). **G)** Heatmap showing the activation of core BX genes in SHWs after D-integration. Expected, expression level before polyploidization; observed, expression level after polyploidization. Expression level is given as log_2_ (tpm + 1) transformed. **H)** Relative contents of core metabolites in BX pathway of hexaploid SHWs and its polyploidization parents. Content level is calculated by log_10_ transformed.

Considering the predominance of glycosylated flavonoids in SHWs, we conducted an *in-vitro* assay to validate the biochemical function of two *O*-glucosyltransferases encoded by D homoeolog genes namely *TraesCS2D01G069100* and *TraesCS2D01G078700* (**Fig. 6C**). The purified protein encoded by *TraesCS2D01G069100* showed glucosyltransferase activity with apigenin, luteolin, chrysoeriol and tricin, with the highest activity observed on provision of apigenin and luteolin as substrates (**Fig. 6D**). Similarly, the purified protein encoded by *TraesCS2D01G078700* showed activity of glycosyl transfer to apigenin and luteolin (**Supplementary Fig. S17**). Additionally, we observed the activation of several putative flavonoid related genes following DD genome integration (**Fig. 6E**). These findings collectively demonstrate that the activation of the flavonoid pathway in allohexaploid wheat is attributed to combined effects of the introduction of novel functional genes via the DD genome integration and upregulating gene expression in the A and B subgenomes.

It is remarkable that lots of benzoxazinoids (BXs) belong to the ABD dominant category, in which metabolites showed highest content over tetraploid and diploid parents in ternary plot analysis (**Fig. 5, D to F; Supplementary Data Set 24**). BXs are a class of indole-derived plant metabolites that are most specially present in Poaceae plants and some dicots, such as wheat, maize and rye, and showed function in plant biotic, abiotic resistance and allelopathic phenomenon (Handrick et al. 2016; Li et al. 2018b; Zhang et al. 2021a; Stahl 2022). We have identified 18 metabolites belong to Bx pathway (**Fig. 6F**; **Supplementary Fig. S18**), 13 (72.2%), 9 (50%) and three (16.7%) of which were ABD dominant, accounting for 27.1%, 16.7% and 60% in ABD dominant category, in SHW3, SHW4 and SHW5, respectively (**Fig. 5, D to F**). The BX biosynthesis pathway starts from indole- 3-glycerol phosphate, which was hydrolyzed to indole by an indole -glycerol phosphate aldolase (Bx1). Indole was then further converted by four cytochrome P450 enzymes (Bx2-Bx5) (Nomura et al. 2005) and one UDP-glucosyltransferase (UGT, Bx8, Bx9) to form HBOA (2-hydroxy-3,4- dihydro-2H-1,4-benzoxazin-3-one), DIBOA [2,4-Dihydroxy-2H-1,4-benzoxazin-3(4H)-one] and DIBOA-Glc (Nomura et al. 2005; Sue et al. 2011), which were finally converted to other derivatives by Bx6, Bx7, Bx10 (Li et al. 2018b; Bakera and Rakoczy-Trojanowska 2020; Shavit et al. 2022) and other unknown enzymes (**Fig. 6F**; **Supplementary Fig. S18A**). BXs are stored as inactive and stable glycosides in the vacuole where they wait their release by conversion to aglucones by β- glucosidases (Wouters et al. 2016; Niculaes et al. 2018).

We noticed that most of the wheat BX related genes showed increased expression levels following polyploidization, especially *TaBx3*, *TaBx4*, *TaBx5* and *Taglu* (**Fig. 6G**), which resulted obvious higher content of almost all metabolites in the BX pathway in SHWs compared with either their tetraploid or diploid parents (**Fig. 6H**; **Supplementary Fig. S18B**). It is suggested that B subgenome homoeologs of wheat BX genes mainly contributed to BXs synthesis, which in turn was contributed by the different progenitors of the subgenomes (Nomura et al. 2005; Sue et al. 2011). The higher expression level of B homoeologs of *TaBx3*, *TaBx4*, *TaBx5* and *Taglu* (*Taglu1a*, *Taglu1b*) than A and D homoeologs was also demonstrated in our study (**Fig. 6G**). Unlike the flavonoid pathway, BX genes from the D genome may contribute little to BX biosynthesis in hexaploid wheat considering the low or even lack of expression of D homoeologs. Our results showing the activation of BX genes, especially B homoeologs, by DD-genome integrated polyploidization events (**Fig. 6G**). Therefore, there may be unknown activators that are introduced from the DD genome integration event which increase transcriptional efficiency and enhanced their expression level, resulting in an overdominance phenomenon of BX biosynthesis in the hexaploidy wheat.

## Discussion

Allopolyploidization is caused by the merging of two or more genomes from divergent parental species in a common nucleus. This sudden perturbation results in broad genetic, epigenetic and gene-expression alterations (McClintock 1984; Li et al. 2011; Matsuoka et al. 2014). Bread wheat is a typical allohexaploid plant that experienced two rounds of polyploidization events during its evolution (International-Wheat-Genome-Sequencing-Consortium 2014, 2018). Integration of the DD genome represent the second polyploidization event and played a vital role in allowing wheat adaptation to diverse environmental conditions and cultivation areas (Dubcovsky and Dvorak 2007; Luo et al. 2017). Here, we comprehensively reveal the impact of DD genome integration from various aspects including gene expression, epigenome organization, chromatin interaction and metabolites.

It has been reported that a large number of genes from the newly introduced D genome were suppressed during wheat polyploidization (Vasudevan et al. 2023), which is in line with the observation here (**Fig. 1, C** and **F; Supplementary Fig. S13**). Interestingly, the reshaping occurred not only in the newly introduced D subgenome, but also affect the original A and B subgenomes, spanning multiple levels including transcriptome (**Fig. 1C**), epigenome (**Fig. 1H**), chromatin interactome (**Fig. 2G; Supplementary Fig. S12E and S13**) and metabolome (**Fig. 6, E and G**), although the D subgenome admittedly underwent the greatest alteration during the process of polyploidization, suggesting extensive interactions occurred among subgenomes during polyploidization. Indeed, the chromatin interactome data demonstrated massive newly built interchromosomal interactions between A or B subgenome with the integrated D subgenome (**Fig. 2G**), which enhanced the interactions and connections between subgenomes and largely improved genomic robustness, genetic diversity and plasticity of gene transcription regulation in hexaploid wheat.

Hexaploid wheat rapidly surpassed its tetraploid ancestor species within a short period of time and became the most widely planted crop globally, accounting for more than 95% of all wheat-growing areas, likely due to the improved environmental adaptability in hexaploid wheat (Dubcovsky and Dvorak 2007). Both of our current study and other reports showed that hexaploid wheat is more tolerant to UV-B light, salt stress, nitrogen deficiency and zinc deficiency than tetraploid or diploid materials (**Fig. 4, D** to **H**; **Supplementary Fig. S14**; Cakmak et al. 1999; Yang et al. 2014; Yang et al. 2018). However, the mechanism of how adaptability increased in hexaploidy is yet to be clarified. Our studies revealed that the enhanced adaptability of hexaploidy wheat is the result of a coordinated action at various levels in wheat genome. The transcriptome analysis demonstrated the upregulation of many environmental adaptation and stress-response-related genes during DD genome integration, both in the A, B and D subgenome (**Fig. 1D; Supplementary Fig. S5A**). Epigenomic results suggested the enrichment of active histone modifications on different stress and environment response genes (**Fig. 1I**). Moreover, 3D interactome analysis revealed that lots of environmental adaptation related genes were connected with more interaction loops following polyploidization (**Fig. 2J**). Furthermore, substantial enrichment of secondary metabolites related to stress responses have been observed in metabolome analysis **(Fig. 5 and 6**). We speculated that a tight connection of these adaptation genes in the wheat genome mediated by the DD- integration may facilitate their high-efficiency co-transcription. This hypothesis was supported by the results that significant more loops were connected with RLKs and chitinase encoding genes and with genes reported to be associated with stresses such as high alkalinity, high salinity, UV-B exposure, cold, low nitrogen, and flavonoid metabolism, subsequently activating transcription of these tightly connected genes (**Fig. 3 and 4)**.

We further assessed the common metabolic features of polyploidization in our studies. An increased content of alkaloids and a decrease of carbohydrates was found in SHWs following polyploidization events (**Fig. 5, B to I**). Whilst secondary metabolites were not measured in the previous study using metabolomes to evaluate changes upon domestication of tetraploid durum wheat carbohydrates levels were largely unaltered (Beleggia et al. 2016). This comparison giving further support to the thesis that the decreased carbohydrates are the result of the integration of the DD genome. Whether the decrease in carbohydrates is the result of increased carbon partitioning to alkaloids and/or phenylpropanoids will however require further, more detailed investigation. However, it is interesting to note that the metabolic changes during domestication appear to be largely species specific. A recent review describing the consequences of domestication of the metabolome on rice, maize and wheat as well as tomato melon and tea concluded this (Alseekh et al. 2021a). Moreover, data in subsequent studies in jujube, pepper and peach (Cao et al. 2022), confirm this. In addition, our results strongly suggest that the changes caused by polyploidization are directional and reproducible both in transcriptome and epigenome. Indeed, the detailed direction and common features of various aspects have comprehensively and systemically demonstrated in our study for the first time.

We found the flavonoid pathway, which plays important roles in oxidation and UV resistance (Fang et al. 2019), was activated in the SHWs compared to their tetraploid parent (**Fig. 6, A and B**). This phenomenon possibly resulted from the addition of putative flavonoid synthesis genes located on the D subgenome (**Fig. 6C**). Indeed, the catalytic activities of two such glucosyltransferases acting in the flavonoid pathway encoded by D subgenome genes were confirmed in our present study (**Fig. 6D**; **Supplementary Fig. S17**), with our results suggesting the integration of these genes from the DD genome to allohexaploid wheat may enhance the content of flavonoid glycosides. This is however unlikely to be the only reason as we also found activation of candidate flavonoid genes in the A, B and D subgenomes following polyploidization (**Fig. 6E**), which may also contribute to the enhanced flavonoid content. In addition, we found an enhanced transcription of BX biosynthesis genes, especially the B homoeologs, accompanied with activation of entire BX biosynthesis pathway in hexaploid wheat after polyploidization (**Fig. 6, G and H; Supplementary Fig. S18**). BXs are widely reported to function in environmental adaptability (Handrick et al. 2016; Li et al. 2018b; Zhang et al. 2021a; Batyrshina et al. 2022). Considering the extremely low or lack of D homoeologs expression of key BX genes, we speculate a putative transcription activator been introduced to hexaploid wheat through polyploidization to promote the transcription of A, B subgenome-located BX genes, resulting an over-parental expression level of genes involved in BX pathway. However, whether both contribute and if so, what is the molecular hierarchy of these events requires to be clarified in future studies. Taking above mentioned results together, changed gene transcription network and metabolome mediated by DD integration facilitated the worldwide spread of allohexaploid wheat and subsequently rendered the plant as an indispensable global crop within a relatively short frame of time.

In conclusion, here we provide a comprehensive and systematic multi-omics landscape resource including transcriptome, epigenome, 3D chromatin interactions and metabolome to understand the influence of DD genome integration-based polyploidization events. We illuminate common and reproducible directions of wheat polyploidization. Through integrated analysis of transcriptome, 3D interactome and metabolome atlases, we revealed that polyploidization-mediated dynamics in gene transcription and metabolites increased adaptability of allohexaploid wheat to various environment via coordinated effects among subgenomes and the upregulation of protective small molecules. Whilst our study provides considerable insight into this aspect of wheat evolution, we believe that the datasets produced will additionally enable researchers to probe a large number of other important questions in wheat biology.

## Methods

### Plant materials, growth conditions and stress treatments

Three synthetic hexaploid wheat (SHW3, SHW4, SHW5; AABBDD), their related tetraploid durum wheat (AABB) and diploid *Ae. tauschii* (DD), and the hexaploid bread wheat Aikang 58 were used in this study. Detail information of these materials is provided in **Supplementary Data Set 1**.

Materials used for stress treatments were soil grown in plastic pots under normal conditions for 18 days (three-leaves stage) with 16 h light and 8 h dark photoperiod at 21°C ± 1°C in greenhouse. Each pot contained nine seedlings and served as a biological replicate. For alkaline stress, 100 mM mixed alkali (NaHCO_3_:Na_2_CO_3_ with a molar ratio of 5:1, 83.3 mM for NaHCO_3_ and 16.7 mM for Na_2_CO_3_ respectively; PH = 9.5) was used to treat the 18-days seedlings for 28-33 days. For salt treatment, 250 mM NaCl were used to irrigate these seedlings for 28 days. Another group of these materials were continued to grow at normal conditions irrigating with water for control. Subsequently, 18-days seedlings grown under normal conditions were transferred to a UV- B chamber (TL8W/302 nm narrowband UV-B tube) without white light. The UV-B intensity was 12.8 μW cm^-2^ measured using a UV radiometer. After 9 days UV-B stress, all the materials were transferred back to normal light conditions for recovery. Two weeks later, the growth and survival status of these plants were recorded.

For multi-omics analysis, materials including SHWs, and its tetraploid and diploid parents were soil grown in greenhouse with 12 h light and 12 h dark cycle at constant temperature (16°C ± 1°C). The top fully expanded leaves of 21-days seedlings were harvested without crosslink and stored at -80°C for RNA sequencing (RNA-seq) and metabolome analysis. Leaf samples for ChIP-seq and ChIA-PET experiments were cut into pieces for crosslinking after been harvested immediately. Samples for ATAC-seq experiments were used fresh leaves without crosslinking.

### RNA-seq library preparation

Total RNA were isolated from wheat leaves using TRIzol reagent (Invitrogen) according to protocol (Fang et al. 2018). RNA libraries were constructed using MGIEasy RNA Library Prep Kit. RNA sequencings were performed using an MGI DNBseq-T7 (paired-end 150 bp reads) system. Three biological replicates of RNA-seq data were obtained from each material.

### ChIP-seq library preparation

ChIP-seq experiments were performed according to published protocols (Kaufmann et al. 2010; Zhao et al. 2020) with modifications. Wheat leaves were cut into pieces then cross-linked with 1% formaldehyde in MC buffer (Kaufmann et al. 2010) for 20 min and quenched using 1.25 M glycine (final concentration was 0.125 M). About 0.2 g cross-linked tissues were grind into fine power in liquid nitrogen for each ChIP assay. Then, 300 μL Buffer S were added to lyse for 30 min at 4°C and 700 μL Buffer F for 15 min with rotation at 4°C. Chromatin were fragmented to 200-700 bp by sonication using Covaris S220 sonicator (peak power 150, percentage 6%, cycles per burst 200) for 10 min. After two rounds of centrifugation at maximum speed for 10 min at 4°C, supernatant containing sonicated chromatin were transferred to new tube for pre-clearing rotation with 20 μL Dynabeads^TM^ protein A (Invitrogen, 10002D) at 4°C for 1 h. Then another 40 μL Dynabeads^TM^ protein A and 5 μL antibodies respectively against H3K4me3 (ABclonal, A2357), H3K9ac (Cell Signaling Technology, 9649S), H3K27me3 (Cell Signaling Technology, 9733S) were added to the chromatin supernatant for immunoprecipitation (IP) with rotation at 4°C overnight. After washed with low-salt ChIP buffer, high-salt ChIP buffer, ChIP wash/LiCl buffer and TE buffer in order, the protein-DNA complexes were eluted from Dynabeads^TM^ protein A using ChIP elution buffer by two rounds at 65°C for 15 min with agitation at 900 rpm. 15 μL 5 M NaCl and 13 μL proteinase K (20 mg/mL) were added to the eluate for reverse crosslinking at 65°C for 8 h. ChIPed DNA were precipitated by pre-cooled ethanol and purified using QIAquick PCR Purification Kit (QIAGEN, 28106), then resolved with 25 μL Buffer EB. ChIP-seq libraries were prepared using ThruPLEX DNA-seq 48S Kit (Takara, R400427) following the manufacturer and sequenced using Illumina NovaSeq 6000 system (paired-end 150 bp reads).

### ATAC-seq libraries preparation

To construct ATAC-seq libraries, fresh tissues were cut into pieces and chopped into homogenate in pre-chilled D-Sorbitol buffer (0.6 M D-sorbitol, 20 mM MES, 20 mM KCl, 10 mM MgCl_2_, 1 × PMSF, 0.2% Triton X-100, 1 mM β-mercaptoethanol, 0.1 mg/mL BSA, 1 × protease inhibitor) (Tu et al. 2020). Then, the homogenate was filtered through 30 μm cell strainer twice. After stained with 4’,6-diamidino-2-phenylindole (DAPI), crude nuclei were loaded into a flow cytometer (BD FacsAria II SORP). A total of 50,000 nuclei were sorted and collected in pre-chilled D-Sorbitol buffer, then the nuclei were spun down at 1000 g for 10 min at 4°C. After supernatant removed carefully, nuclei were mixed with 1 × TD buffer (10 mM Tris-HCl, 5 mM MgCl_2_, 10% DMF) and 1.5 μL TTE Mix (Vazyme, TD501) and incubated at 37°C for 30 min with mixing gently every 5 min. Reactions were stopped using Buffer PB and DNA were purified by MinElute PCR Purification Kit (QIAGEN, 28006). ATAC-seq libraries were prepared with NEBNext^®^ High-Fidelity 2X PCR Master Mix (NEB, M0541L) and TruePrep^®^ Index Kit V2 for Illumina (Vazyme, TD202). Libraries were sequenced using Illumina NovaSeq 6000 system (paired-end 150 bp reads).

### Long-read ChIA-PET library preparation

H3K4me3- associated long-read ChIA-PET library construction were performed according to published protocol with minor modifications (Tang et al. 2015; Li et al. 2017; Zhao et al. 2019). Briefly, fresh wheat leaves were cut into pieces and treated with 1.5 mM Ethylene glycol bis (succinimidylsuccinate) (EGS, Thermo Scientific, 21565) under vacuum for 20 min crosslinking. Then formaldehyde (Thermo Scientific, 28906) was added to 1% final concentration for another 20 min crosslinking. One tenth volume of 1.25 M glycine were added to stop the crosslinking reaction and wished three times with ddH_2_O. Cross-linked tissues were stored at -80°C until use. About 4.5 g sample was used for each ChIA-PET experiment. After ground into fine powder in liquid nitrogen, tissues were mixed thoroughly with 6.75 mL 1% SDS FA nuclear lysis buffer and incubated at 4°C for 30 min with rotation and 27 mL SDS-free FA lysis buffer for another 15 min incubation at 4°C. The solution was sonicated to fragment chromatin into 1-3 kb using a BioRuptor (Diagenode) under high level intensity (30 sec ON and 50 sec OFF, 20 cycles). After centrifugation at 12000 rpm for 10 min at 4°C, the supernatant was transferred to new tube for next immunoprecipitation. About 800 μL Dynabeads^TM^ protein A were mixed with 100 μL H3K4me3 antibody (ABclonal, A2357) and incubated at 4°C for 8 h to generate antibody-coated beads. Then, the fragmented chromatin was incubated with antibody-coated beads at 4°C overnight with rotation. The ChIPed beads were orderly washed with 5 mL 0.1% SDS FA cell lysis buffer three times, 5 mL High salt ChIP buffer twice, 5 mL ChIP wash buffer once and 5 mL TE buffer once. After end-repairing and A-tailing, biotinylated bridge linker was used to perform proximity ligation of ChIP DNA on the beads. The proximity ligated ChIP DNA were reverse cross-linked using proteinase K and extracted by phenol-chloroform alcohol. ChIA-PET libraries were constructed using Tn5 transposase (TruePrep DNA Library Prep Kit V2 for Illumina, Vazyme, TD501) and sequenced using Illumina NovaSeq 6000 system (paired-end 150 bp reads).

### Metabolic profiling and metabolite annotation

Metabolite extraction and detection was performed according to previous research (Chen et al. 2013). The freeze-dried wheat leaves were pulverized by a mixer mill at 30 Hz for 1 min. 100 mg powder was used to extract metabolites for each biological replicates by adding 1 mL pre-cooled 70% methanol (containing 0.1 mg/L acyclovir as internal standard). After mixing thoroughly, the homogenate was incubated in an ultra-sonication bath for 15 min at 4°C with vortexing every 3 min or incubated at 4 °C with rotation overnight, then followed centrifugation at 12000 *g* at 4°C for 10 min. The supernatant was filtered through a 0.22 μm filter membrane (ANPEL, SCAA-104). The final filtrates were analyzed using liquid chromatography-electrospray ionization- tandem mass spectrometry (LC-ESI-MS/MS) system. The MS2T library was established using a stepwise multiple ion monitoring-enhanced product ion according to previous published protocols (Chen et al. 2013). To facilitate annotation of metabolites, the detailed information contained in metabolite signals were compared with metabolite databases and available commercial standards. The scheduled multiple reaction monitoring (MRM) method was used to quantified the relative contents of the 839 metabolites identified in this study as previously described (Chen et al. 2014). The scheduled MRM algorithm was used with an MRM detection window of 90 sec and a target scan time of 1 sec using Analyst 1.5 software. Metabolite data are reported following set reporting standards (Fernie et al. 2011; Alseekh et al. 2021b).

### Enzyme activity assay

The full-length cDNA of *TraesCS2D01G078700* and *TraesCS2D01G069100* were amplified using specific primers (**Supplementary Data Set 25**) and respectively cloned into expression vector pGEX-6p-1 that posse a GST tag at N terminus. Recombinant vectors were transformed into *Escherichia coli* expression strain BL21 (DE3), respectively. Expressed GST-tagged proteins were harvested from lysed *E. coli* cells and purified using Glutathione Sepharose 4B (GE Healthcare). The glucosyltransferase activities of recombinant GST-tagged proteins were analyzed according to published method (Chen et al. 2020a). In brief, purified proteins were added into a total of 10 μL reaction mixture containing 200 mM Tris-HCl (PH7.5), 10 mM MgCl_2_, 0.75 mM UDP-glucose and 0.1 mM substrate (apigenin, luteolin, chrysoeriol and tricin respectively) and incubated at 37°C for 30 min, then the reactions were stopped by addition of 30 μL methanol. The generated glycosylation products were detected by LC-MS/MS analysis. Expressed protein from pGEX-6p-1 empty vector was used as negative controls.

### RNA-seq data analysis

RNA-seq reads were aligned to the Chinese Spring RefSeqv1.0 reference genome (International- Wheat-Genome-Sequencing-Consortium 2018) using STAR (Dobin et al. 2012) (version 2.7.1a). Materials of different ploidy were individually mapped to the corresponding genomes, the tetraploid datasets of AABB were only aligned to A and B subgenome transcripts and the diploid of DD was only aligned to D subgenome transcripts, the diploid of DD was only aligned to the D subgenome sequences and the hexaploidy of AABBDD was aligned to the A, B, and D subgenome transcripts. The expression of annotated genes was measured by RSEM (Li and Dewey 2011) (version 1.2.22) and normalized with transcripts per million (TPM). We considered the gene with TPM > 0.5 expressed. For gene quantification comparison, we referred to published research (Ramírez-González et al. 2018) and combined the TPM matrices of AABB and DD in a 2:1 ratio and then normalized them to 10^6^ as SHW. We consider a gene as differentially expressed gene (DEG) when: 1. TPM in expected and observed > 0.5, fold change >2 and FDR <0.05; or 2. TPM in expected = 0 and observed > 0.5, or expected > 0.5 and observed = 0; or 3. TPM in 0 < expected < 0.5 and observed > 1, or expected > 1 and 0 < observed < 0.5. For expression breadth analysis, we downloaded the transcription data of 23 different tissues from published research (Ramírez-González et al. 2018) (**Supplementary Data Set 12**, 3 replicates were included), and the gene with TPM > 1.5 were considered as expressed in this tissue.

### Homoeolog expression bias analysis

We first downloaded gene triads sets with A: B: D = 1:1:1 correspondence in three homologous subgenomes (Ramírez-González et al. 2018). We extracted these gene TPM values from all genes’ TPM matrix generated above and normalized the TPM for each gene within the triad according to published method (Ramírez-González et al. 2018). The TPM sum of the gene triad below 0.5 was excluded from the downstream analysis. Then we calculated the relative contributions of these genes in the respective triad and calculated the Euclidean distance to the ideal normalized expression bias table [Table S37 in ref. (Ramírez-González et al. 2018)] with the rdist function from rdist R package. We group each triad into the same category as the one with the shortest distance from the seven ideal categories.

### GO analysis

The gene ontology (GO) enrichment analyses were performed on the Triticeae-Gene Tribe database (Chen et al. 2020b) (http://wheat.cau.edu.cn/TGT/). The GO term with a Bonferroni corrected *P*- value less than 0.05 we considered as significant.

### ATAC-seq data analysis

For the pre-processing of ATAC-seq data, we follow the workflow of ChIP-Hub (Fu et al. 2022). The raw reads were first trimmed by Trimmomatic (Bolger et al. 2014) (version 0.36) to remove sequencing adapters. The trimmed reads were aligned to the Chinese Spring RefSeqv1.0 reference genome (International-Wheat-Genome-Sequencing-Consortium 2018) using Bowtie2 (Langmead and Salzberg 2012) with the following parameters “-q—no-unal—threads 8—sensitive”. All mapped to mitochondrial and chloroplast DNA reads were removed. After sorting mapped reads with SAMtools (Li and Durbin 2009) (version 0.1.19), we only used properly paired reads with high mapping quality (MAPQ score > 30) for the subsequent analysis. The PCR duplicates were removed using the MarkDuplicates function from Picard tools (version 2.60; http://broadinstitute.github.io/picard/). MACS2 (Zhang et al. 2008) (version 2.1.0) was used to call peaks. Using the “callpeak” function in MACS2 with the following parameters: “--call-summits -f BAM --nomodel --mfold 2 20 --qvalue 0.01”. The “-shift” used in the model was determined by the analysis of cross-correlation scores using the phantompeakqualtools package (https://code.google.com/archive/p/phantompeakqualtools/). The “--gsize” was set to the corresponding subgenome size. The final peaks were reproducible across pseudo-replicates and true replicates with an IDR (Irreproducibility Discovery Rate) < 0.05. For quantification, the Tn5 insertion positions were first determined as the start sites of reads adjusted by the rule of “forward strand +4 bp, negative strand −5 bp” (Buenrostro et al. 2013). The peak sets were merged by the merge function from BEDTools (Quinlan and Hall 2010) (version 2.30.0), the number of Tn5 cuts in each peak was counted, and the sum is normalized to 10^6^ by dividing by the length of the peaks.

### ChIP-seq data analysis

We used the ChIP-Hub (Fu et al. 2022) analysis workflow to process the ChIP-seq data. The sequencing data trimming, reads alignment, filtering, etc. were all processed in the same manner as the ATAC-seq. The peak calling was also similar to that of ATAC-seq, except that the input sample was added here as a control.

### Differential peak analysis

We merged the peaks of H3K9ac, H3K27me3, H3K4me3, and ATAC-seq separately. For the histone modification data, we counted the number of reads in the corresponding peak and divide it by the length of the peak, and finally normalized it to 10^6^. For ATAC-seq, we counted the insertion position of Tn5 as described above. Then we used the Shannon entropy method to screen differential peaks. We calculated the entropy score for each peak using the philentropy (https://github.com/drostlab/philentropy) R package and considered the entropy value less than 1.6 as differential peaks. Finally, we performed Z-score transformation by row for visualization. For the target genes of these differential peaks, we assigned these peaks to the nearest TSSs according to their summit positions using BEDTools (Quinlan and Hall 2010) closest function to identify target genes. The GO enrichment of the target gene was performed as described above.

### Pseudo-bam generated and genome tracks visualization

To compare the differences reflected by the various data during the polyploidization, we synthesized pseudo-bam by mapping reads to the A, B, D subgenomes in a 1:1:1 ratio. Then we used the bamCoverage function from deepTools (Ramírez et al. 2014) (version 3.5.1) package with the following parameters “--normalizeUsing RPKM --binSize 5” to generate bigWig file. We used the WashU Epigenome Browser (Li et al. 2022a) to visualize signal tracks with bigWig files.

### ChIA-PET data analysis

The ChIA-PET data were analyzed using updated ChIA-PET Tool software package for automatic processing of sequence data (Li et al. 2010), including linker filtering, read mapping to reference genome (International-Wheat-Genome-Sequencing-Consortium 2018) using BWA, redundancy removal, identification of protein binding sites and chromatin interactions. Mapped PETs were classified to self-ligation, inter-ligation and other PETs according to genomic span of a PET. Obtained inter-ligation PETs (genomic span > 8 kb) were used to generate raw contact frequency matrix and other future analysis. Considering the large genome size and intergenic regions of wheat, we chose ChIP-seq peak summit ± 3 kb as the given anchors to call clusters with removal of overlapping with adjacent anchor region. We defined a loop as PPI interaction when the two ChIP-seq peak summit all located in promoter regions, and as PDI interaction when one peak summit located in promoter regions and another peak summit located in distal regions (regions outside promoter and gene body). We also defined a loop as intrachromosomal loop when the two anchors located in the same chromosome and as interchromosomal loop when the two anchors located in different chromosomes.

### Network constructions

The interaction networks were constructed through two or three hops of interactions (including PPI and PDI) originating from these selected core DEGs or adaptive genes. Nodes, including anchor genes and distal elements, were connected on the basis of chromatin interactions present in the ChIA-PET libraries from SHW3 (after polyploidization) and its tetraploid, diploid parents (before polyploidization) respectively. The connectivity networks were visualized and modularized using Gephi (Bastian et al. 2009).

### Data availability

The RNA-seq, ATAC-seq, ChIP-seq and ChIA-PET data involved in this study are available at the NCBI BioProject (PRJNA939846).

### Code availability

The data analysis code has been stored as Jupyter notebooks in the GitHub repository (https://github.com/tzhu-bio/Wheat_Polyploidization_Analysis).

## Acknowledgments

We thank Prof. Guoliang Li, Dr. Qiangwei Zhou and Dr. Xingyu Huang (Huazhong Agricultural University) for their valuable assistance in ChIA-PET data analysis. This work was supported by grants from National Key Research and Development Program of China (2023YFF1001302), Biological Breeding-National Science and Technology Major Project (2023ZD0407105), the National Key Laboratory of Crop Genetic Improvement Self-Research Program (ZW19A0201), the National Natural Science Foundation of China (32202248 and 32070656) and the China Postdoctoral Science Foundation (2019M662665).

## Author Contributions

W.Y. conceived the study with input from D.C. and A.R.F.; Y.L. conducted most of experiments and generated the data with assistance from W.C., X.W., C.C., S.B., W.O., Q.L., H.M. and X.L.; D.C., T.Z. and Y.L. performed data analysis with support from X.Z., C.H., W.C. X.H. and S.B.; J.W. and M.L. performed enzyme activity assay; M.K. and C.L. provided the materials; H.S. performed cytology karyotyping analysis; Y.L. and T.Z. drafted the manuscript; W.Y., D.C., A.R.F. and K.K. discussed and revised the manuscript.

## References

Alseekh, S., Scossa, F., Wen, W., Luo, J., Yan, J., Beleggia, R., Klee, H.J., Huang, S., Papa, R., and Fernie, A.R. (2021a). Domestication of Crop Metabolomes: Desired and Unintended Consequences. Trends Plant Sci 26, 650–661.

Alseekh, S., Aharoni, A., Brotman, Y., Contrepois, K., D’Auria, J., Ewald, J. J. C.E., Fraser, P.D., Giavalisco, P., Hall, R.D., et al. (2021b). Mass spectrometry-based metabolomics: a guide for annotation, quantification and best reporting practices. Nat Methods 18, 747–756.

Avni, R., Lux, T., Minz-Dub, A., Millet, E., Sela, H., Distelfeld, A., Deek, J., Yu, G., Steuernagel, B., Pozniak, C., et al. (2022). Genome sequences of three Aegilops species of the section Sitopsis reveal phylogenetic relationships and provide resources for wheat improvement. Plant J 110, 179–192.

Bakera, B., and Rakoczy-Trojanowska, M. (2020). Isolation and structural analysis of the Bx6 and Bx7 genes controlling the biosynthesis of benzoxazinoids in rye (Secale cereale L.). Acta Physiol Plant 42, 56.

Bastian, M., Heymann, S., and Jacomy, M. (2009). Gephi: An Open Source Software for Exploring and Manipulating Networks. Proceedings of the International AAAI Conference on Web and Social Media 3, 361–362.

Batyrshina, Z.S., Shavit, R., Yaakov, B., Bocobza, S., and Tzin, V. (2022). The transcription factor TaMYB31 regulates the benzoxazinoid biosynthetic pathway in wheat. J Exp Bot 73, 5634–5649.

Beleggia, R., Rau, D., Laido, G., Platani, C., Nigro, F., Fragasso, M., De Vita, P., Scossa, F., Fernie, A.R., Nikoloski, Z., et al. (2016). Evolutionary metabolomics reveals domestication-associated changes in tetraploid wheat kernels. Mol Biol Evol 33, 1740–1753.

Bolger, A.M., Lohse, M., and Usadel, B. (2014). Trimmomatic: a flexible trimmer for Illumina sequence data. Bioinformatics 30, 2114–2120.

Buenrostro, J.D., Giresi, P.G., Zaba, L.C., Chang, H.Y., and Greenleaf, W.J. (2013). Transposition of native chromatin for fast and sensitive epigenomic profiling of open chromatin, DNA-binding proteins and nucleosome position. Nat Methods 10, 1213–1218.

Cakmak, I., Tolay, I., Ozdemir, A., Ozkan, H., Ozturk, L., and Kling, C.I. (1999). Differences in Zinc Efficiency among and within Diploid, Tetraploid and Hexaploid Wheats. Ann Bot 84, 163–171.

Cao, K., Wang, B., Fang, W., Zhu, G., Chen, C., Wang, X., Li, Y., Wu, J., Tang, T., Fei, Z., et al. (2022). Combined nature and human selections reshaped peach fruit metabolome. Genome Biol 23, 146.

Cavalet-Giorsa, E., Gonzalez-Munoz, A., Athiyannan, N., Holden, S., Salhi, A., Gardener, C., Quiroz-Chavez, J., Rustamova, S.M., Elkot, A.F., Patpour, M., et al. (2024). Origin and evolution of the bread wheat D genome. Nature.

Chen, J., Hu, X., Shi, T., Yin, H., Sun, D., Hao, Y., Xia, X., Luo, J., Fernie, A.R., He, Z., et al. (2020a). Metabolite-based genome-wide association study enables dissection of the flavonoid decoration pathway of wheat kernels. Plant Biotechnol J 18, 1722–1735.

Chen, J., Wang, J., Chen, W., Sun, W., Peng, M., Yuan, Z., Shen, S., Xie, K., Jin, C., Sun, Y., et al. (2018). Metabolome analysis of multi-connected biparental chromosome segment substitution line populations. Plant Physiol 178, 612–625.

Chen, W., Gong, L., Guo, Z., Wang, W., Zhang, H., Liu, X., Yu, S., Xiong, L., and Luo, J. (2013). A novel integrated method for large-scale detection, identification, and quantification of widely targeted metabolites: application in the study of rice metabolomics. Mol Plant 6, 1769–1780.

Chen, W., Gao, Y., Xie, W., Gong, L., Lu, K., Wang, W., Li, Y., Liu, X., Zhang, H., Dong, H., et al. (2014). Genome-wide association analyses provide genetic and biochemical insights into natural variation in rice metabolism. Nat Genet 46, 714–721.

Chen, Y., Song, W., Xie, X., Wang, Z., Guan, P., Peng, H., Jiao, Y., Ni, Z., Sun, Q., and Guo, W. (2020b). A Collinearity-incorporating Homology Inference Strategy for Connecting Emerging Assemblies in Triticeae Tribe as a Pilot Practice in the Plant Pangenomic Era. Mol Plant 13, 1694–1708.

Chen, Z.J. (2007). Genetic and epigenetic mechanisms for gene expression and phenotypic variation in plant polyploids. Annu Rev Plant Biol 58, 377–406.

Concia, L., Veluchamy, A., Ramirez-Prado, J.S., Martin-Ramirez, A., Huang, Y., Perez, M., Domenichini, S., Rodriguez Granados, N.Y., Kim, S., Blein, T., et al. (2020). Wheat chromatin architecture is organized in genome territories and transcription factories. Genome Biol 21, 104.

Cui, M., Li, Y., Li, J., Yin, F., Chen, X., Qin, L., Wei, L., Xia, G., and Liu, S. (2023). Ca2+-dependent TaCCD1 cooperates with TaSAUR215 to enhance plasma membrane H+-ATPase activity and alkali stress tolerance by inhibiting PP2C-mediated dephosphorylation of TaHA2 in wheat. Mol Plant 16, 571–587.

Deng, L., Gao, B., Zhao, L., Zhang, Y., Zhang, Q., Guo, M., Yang, Y., Wang, S., Xie, L., Lou, H., et al. (2022). Diurnal RNAPII-tethered chromatin interactions are associated with rhythmic gene expression in rice. Genome Biol 23, 7.

Dobin, A., Davis, C.A., Schlesinger, F., Drenkow, J., Zaleski, C., Jha, S., Batut, P., Chaisson, M., and Gingeras, T.R. (2012). STAR: ultrafast universal RNA-seq aligner. Bioinformatics 29, 15–21.

Dubcovsky, J., and Dvorak, J. (2007). Genome plasticity a key factor in the success of polyploid wheat under domestication. Science 316, 1862–1866.

Fang, C., Fernie, A.R., and Luo, J. (2019). Exploring the diversity of plant metabolism. Trends Plant Sci 24, 83–98.

Fang, Y.J., Shen, J.Q., Ma, S.Q., and Xiong, L.Z. (2018). Extraction of Total RNA from Rice Tissues. Bio-protocol, e1010111.

Fang, Z., and Morrell, P.L. (2016). Domestication: Polyploidy boosts domestication. Nat Plants 2, 16116.

Fasano, C., Diretto, G., Aversano, R., D’Agostino, N., Di Matteo, A., Frusciante, L., Giuliano, G., and Carputo, D. (2016). Transcriptome and metabolome of synthetic Solanum autotetraploids reveal key genomic stress events following polyploidization. New Phytol 210, 1382–1394.

Fernie, A.R., Aharoni, A., Willmitzer, L., Stitt, M., Tohge, T., Kopka, J., Carroll, A.J., Saito, K., Fraser, P.D., and DeLuca, V. (2011). Recommendations for reporting metabolite data. Plant Cell 23, 2477–2482.

Fu, L.-Y., Zhu, T., Zhou, X., Yu, R., He, Z., Zhang, P., Wu, Z., Chen, M., Kaufmann, K., and Chen, D. (2022). ChIP-Hub provides an integrative platform for exploring plant regulome. Nat Commun 13, 3413.

Gaurav, K., Arora, S., Silva, P., Sanchez-Martin, J., Horsnell, R., Gao, L., Brar, G.S., Widrig, V., John Raupp, W., Singh, N., et al. (2021). Population genomic analysis of Aegilops tauschii identifies targets for bread wheat improvement. Nat Biotechnol 40, 422–431.

Grover, A. (2012). Plant chitinases: genetic diversity and physiological roles. Crit Rev Plant Sci 31, 57–73.

Guo, W., Xin, M., Wang, Z., Yao, Y., Hu, Z., Song, W., Yu, K., Chen, Y., Wang, X., Guan, P., et al. (2020). Origin and adaptation to high altitude of Tibetan semi-wild wheat. Nat Commun 11, 5085.

Handrick, V., Robert, C.A., Ahern, K.R., Zhou, S., Machado, R.A., Maag, D., Glauser, G., Fernandez-Penny, F.E., Chandran, J.N., Rodgers-Melnik, E., et al. (2016). Biosynthesis of 8-O-Methylated Benzoxazinoid Defense Compounds in Maize. Plant Cell 28, 1682–1700.

He, F., Pasam, R., Shi, F., Kant, S., Keeble-Gagnere, G., Kay, P., Forrest, K., Fritz, A., Hucl, P., Wiebe, K., et al. (2019). Exome sequencing highlights the role of wild-relative introgression in shaping the adaptive landscape of the wheat genome. Nat Genet 51, 896–904.

Huang, A.C., Jiang, T., Liu, Y.X., Bai, Y.C., Reed, J., Qu, B., Goossens, A., Nutzmann, H.W., Bai, Y., and Osbourn, A. (2019). A specialized metabolic network selectively modulates Arabidopsis root microbiota. Science 364, eaau6389.

International-Wheat-Genome-Sequencing-Consortium. (2014). A chromosome-based draft sequence of the hexaploid bread wheat (Triticum aestivum) genome. Science 345, 1251788.

International-Wheat-Genome-Sequencing-Consortium. (2018). Shifting the limits in wheat research and breeding using a fully annotated reference genome. Science 361, eaar7191.

Jia, J., Xie, Y., Cheng, J., Kong, C., Wang, M., Gao, L., Zhao, F., Guo, J., Wang, K., Li, G., et al. (2021). Homology-mediated inter-chromosomal interactions in hexaploid wheat lead to specific subgenome territories following polyploidization and introgression. Genome Biol 22, 26.

Jia, J., Zhao, G., Li, D., Wang, K., Kong, C., Deng, P., Yan, X., Zhang, X., Lu, Z., Xu, S., et al. (2023). Genome resources for the elite bread wheat cultivar Aikang 58 and mining of elite homeologous haplotypes for accelerating wheat improvement. Mol Plant 16, 1893–1910.

Jordan, K.W., Wang, S., Lun, Y., Gardiner, L.J., MacLachlan, R., Hucl, P., Wiebe, K., Wong, D., Forrest, K.L., Consortium, I., et al. (2015). A haplotype map of allohexaploid wheat reveals distinct patterns of selection on homoeologous genomes. Genome Biol 16, 48.

Kaufmann, K., Muino, J.M., Osteras, M., Farinelli, L., Krajewski, P., and Angenent, G.C. (2010). Chromatin immunoprecipitation (ChIP) of plant transcription factors followed by sequencing (ChIP-SEQ) or hybridization to whole genome arrays (ChIP-CHIP). Nat Protoc 5, 457–472.

Langmead, B., and Salzberg, S.L. (2012). Fast gapped-read alignment with Bowtie 2. Nat Methods 9, 357–359.

Leitch, A.R., and Leitch, I.J. (2008). Genomic plasticity and the diversity of polyploid plants. Science 320, 481–483.

Li, A., Liu, D., Yang, W., Kishii, M., and Mao, L. (2018a). Synthetic hexaploid wheat: yesterday, today, and tomorrow. Engineering 4, 552–558.

Li, A., Liu, D., Wu, J., Zhao, X., Hao, M., Geng, S., Yan, J., Jiang, X., Zhang, L., Wu, J., et al. (2014). mRNA and Small RNA Transcriptomes Reveal Insights into Dynamic Homoeolog Regulation of Allopolyploid Heterosis in Nascent Hexaploid Wheat. Plant Cell 26, 1878–1900.

Li, B., and Dewey, C.N. (2011). RSEM: accurate transcript quantification from RNA-Seq data with or without a reference genome. BMC Bioinformatics 12, 323.

Li, B., Förster, C., Robert, C.A.M., Züst, T., Hu, L., Machado, R.A.R., Berset, J.-D., Handrick, V., Knauer, T., Hensel, G., et al. (2018b). Convergent evolution of a metabolic switch between aphid and caterpillar resistance in cereals. Sci Adv 4, eaat6797.

Li, D., Purushotham, D., Harrison, J.K., Hsu, S., Zhuo, X., Fan, C., Liu, S., Xu, V., Chen, S., Xu, J., et al. (2022a). WashU Epigenome Browser update 2022. Nucleic Acids Res 50, W774–W781.

Li, E., Liu, H., Huang, L., Zhang, X., Dong, X., Song, W., Zhao, H., and Lai, J. (2019a). Long-range interactions between proximal and distal regulatory regions in maize. Nat Commun 10, 2633.

Li, G., Fullwood, M.J., Xu, H., Mulawadi, F.H., Velkov, S., Vega, V., Ariyaratne, P.N., Mohamed, Y.B., Ooi, H.-S., Tennakoon, C., et al. (2010). ChIA-PET tool for comprehensive chromatin interaction analysis with paired-end tag sequencing. Genome Biol 11, R22.

Li, H., and Durbin, R. (2009). Fast and accurate short read alignment with Burrows–Wheeler transform. Bioinformatics 25, 1754–1760.

Li, L.F., Zhang, Z.B., Wang, Z.H., Li, N., Sha, Y., Wang, X.F., Ding, N., Li, Y., Zhao, J., Wu, Y., et al. (2022b). Genome sequences of five Sitopsis species of Aegilops and the origin of polyploid wheat B subgenome. Mol Plant 15, 488–503.

Li, X., Luo, O.J., Wang, P., Zheng, M., Wang, D., Piecuch, E., Zhu, J.J., Tian, S.Z., Tang, Z., Li, G., et al. (2017). Long-read ChIA-PET for base-pair-resolution mapping of haplotype- specific chromatin interactions. Nat Protoc 12, 899–915.

Li, Z., Lu, X., Gao, Y., Liu, S., Tao, M., Xiao, H., Qiao, Y., Zhang, Y., and Luo, J. (2011). Polyploidization and epigenetics. Chinese Sci Bull 56, 245-252.

Li, Z., Wang, M., Lin, K., Xie, Y., Guo, J., Ye, L., Zhuang, Y., Teng, W., Ran, X., Tong, Y., et al. (2019b). The bread wheat epigenomic map reveals distinct chromatin architectural and evolutionary features of functional genetic elements. Genome Biol 20, 139.

Luo, M.C., Gu, Y.Q., Puiu, D., Wang, H., Twardziok, S.O., Deal, K.R., Huo, N., Zhu, T., Wang, L., Wang, Y., et al. (2017). Genome sequence of the progenitor of the wheat D genome Aegilops tauschii. Nature 551, 498–502.

Matsuoka, Y., Takumi, S., and Nasuda, S. (2014). Genetic mechanisms of allopolyploid speciation through hybrid genome doubling: novel insights from wheat (Triticum and Aegilops) studies. Int Rev Cell Mol Biol 309, 199–258.

McClintock, B. (1984). The significance of responses of the genome to challenge. Science 226, 792–801.

Morris, E.R., and Walker, J.C. (2003). Receptor-like protein kinases: the keys to response. Curr Opin Plant Biol 6, 339–342.

Niculaes, C., Abramov, A., Hannemann, L., and Frey, M. (2018). Plant Protection by Benzoxazinoids—Recent Insights into Biosynthesis and Function. Agronomy 8, 143.

Nomura, T., Ishihara, A., Yanagita, R.C., Endo, T.R., and Iwamura, H. (2005). Three genomes differentially contribute to the biosynthesis of benzoxazinones in hexaploid wheat. Proc Natl Acad Sci U S A 102, 16490–16495.

Otto, S.P. (2007). The evolutionary consequences of polyploidy. Cell 131, 452–462.

Pei, H., Teng, W., Gao, L., Gao, H., Ren, X., Liu, Y., Jia, J., Tong, Y., Wang, Y., and Lu, Z. (2022). Low-affinity SPL binding sites contribute to subgenome expression divergence in allohexaploid wheat. Sci China Life Sci 66, 819–834.

Peng, M., Shahzad, R., Gul, A., Subthain, H., Shen, S., Lei, L., Zheng, Z., Zhou, J., Lu, D., Wang, S., et al. (2017). Differentially evolved glucosyltransferases determine natural variation of rice flavone accumulation and UV-tolerance. Nat Commun 8, 1975.

Peng, Y., Xiong, D., Zhao, L., Ouyang, W., Wang, S., Sun, J., Zhang, Q., Guan, P., Xie, L., Li, W., et al. (2019). Chromatin interaction maps reveal genetic regulation for quantitative traits in maize. Nat Commun 10, 2632.

Petersen, G., Seberg, O., Yde, M., and Berthelsen, K. (2006). Phylogenetic relationships of Triticum and Aegilops and evidence for the origin of the A, B, and D genomes of common wheat (Triticum aestivum). Mol Phylogenet Evol 39, 70–82.

Pont, C., Leroy, T., Seidel, M., Tondelli, A., Duchemin, W., Armisen, D., Lang, D., Bustos-Korts, D., Goue, N., Balfourier, F., et al. (2019). Tracing the ancestry of modern bread wheats. Nat Genet 51, 905–911.

Qi, B., Huang, W., Zhu, B., Zhong, X., Guo, J., Zhao, N., Xu, C., Zhang, H., Pang, J., Han, F., et al. (2012). Global transgenerational gene expression dynamics in two newly synthesized allohexaploid wheat (Triticum aestivum) lines. BMC Biol 10, 3.

Quinlan, A.R., and Hall, I.M. (2010). BEDTools: a flexible suite of utilities for comparing genomic features. Bioinformatics 26, 841–842.

Ramírez-González, R.H., Borrill, P., Lang, D., Harrington, S.A., Brinton, J., Venturini, L., Davey, M., Jacobs, J., van Ex, F., Pasha, A., et al. (2018). The transcriptional landscape of polyploid wheat. Science 361, eaar6089.

Ramírez, F., Dündar, F., Diehl, S., Grüning, B.A., and Manke, T. (2014). deepTools: a flexible platform for exploring deep-sequencing data. Nucleic Acids Res 42, W187–191.

Ramsey, J.a., and Schemske, D.W. (1998). Pathways, mechanisms, and rates of polyploid formation in flowering plants. Ann Rev Ecol Syst 29, 467–501.

Ren, Z., Ji, X., Jiao, Z., Luo, Y., Zhang, G.Q., Tao, S., Lei, Z., Zhang, J., Wang, Y., Liu, Z.J., et al. (2020). Functional analysis of a novel C-glycosyltransferase in the orchid Dendrobium catenatum. Hortic Res 7, 111.

Rosyara, U., Kishii, M., Payne, T., Sansaloni, C.P., Singh, R.P., Braun, H.J., and Dreisigacker, S. (2019). Genetic contribution of synthetic hexaploid wheat to CIMMYT’s spring bread wheat breeding germplasm. **Sci Rep** 9, 12355.

Salman-Minkov, A., Sabath, N., and Mayrose, I. (2016). Whole-genome duplication as a key factor in crop domestication. Nat Plants 2, 16115.

Shaked, H., Kashkush, K., Ozkan, H., Feldman, M., and Levy, A.A. (2001). Sequence elimination and cytosine methylation are rapid and reproducible responses of the genome to wide hybridization and allopolyploidy in wheat. Plant Cell 13, 1749-1759.

Shang, Y., Ma, Y., Zhou, Y., Zhang, H., Duan, L., Chen, H., Zeng, J., Zhou, Q., Wang, S., Gu, W., et al. (2014). Biosynthesis, regulation, and domestication of bitterness in cucumber. Science 346, 1084–1088.

Shavit, R., Batyrshina, Z.S., Yaakov, B., Florean, M., Köllner, T.G., and Tzin, V. (2022). The wheat dioxygenase BX6 is involved in the formation of benzoxazinoids in planta and contributes to plant defense against insect herbivores. Plant Sci 316, 111171.

Shumayla, Tyagi, S., and Upadhyay, S.K. (2019). Receptor-like kinases and environmental stress in plants. In Molecular Approaches in Plant Biology and Environmental Challenges, S.P. Singh, S.K. Upadhyay, A. Pandey, and S. Kumar, eds (Singapore: Springer Singapore), pp. 79–102.

Song, Q., and Chen, Z.J. (2015). Epigenetic and developmental regulation in plant polyploids. Curr Opin Plant Biol 24, 101–109.

Stahl, E. (2022). New insights into the transcriptional regulation of benzoxazinoid biosynthesis in wheat. J Exp Bot 73, 5358–5360.

Sue, M., Nakamura, C., and Nomura, T. (2011). Dispersed benzoxazinone gene cluster: molecular characterization and chromosomal localization of glucosyltransferase and glucosidase genes in wheat and rye. Plant Physiol 157, 985–997.

Tang, D., Wang, G., and Zhou, J.M. (2017). Receptor kinases in plant-pathogen interactions: more than pattern recognition. Plant Cell 29, 618–637.

Tang, Z., Luo, O.J., Li, X., Zheng, M., Zhu, J.J., Szalaj, P., Trzaskoma, P., Magalska, A., Wlodarczyk, J., Ruszczycki, B., et al. (2015). CTCF-mediated human 3D genome architecture reveals chromatin topology for transcription. Cell 163, 1611–1627.

Tu, X., Mejia-Guerra, M.K., Valdes Franco, J.A., Tzeng, D., Chu, P.Y., Shen, W., Wei, Y., Dai, X., Li, P., Buckler, E.S., et al. (2020). Reconstructing the maize leaf regulatory network using ChIP-seq data of 104 transcription factors. Nat Commun 11, 5089.

Vaghela, B., Vashi, R., Rajput, K., and Joshi, R. (2022). Plant chitinases and their role in plant defense: A comprehensive review. Enzyme Microb Technol 159, 110055.

Van de Peer, Y., Mizrachi, E., and Marchal, K. (2017). The evolutionary significance of polyploidy. Nat Rev Genet 18, 411–424.

Van de Peer, Y., Ashman, T.L., Soltis, P.S., and Soltis, D.E. (2021). Polyploidy: an evolutionary and ecological force in stressful times. Plant Cell 33, 11–26.

Vasudevan, A., Levesque-Lemay, M., Edwards, T., and Cloutier, S. (2023). Global transcriptome analysis of allopolyploidization reveals large-scale repression of the D-subgenome in synthetic hexaploid wheat. Commun Biol 6, 426.

Walkowiak, S., Gao, L., Monat, C., Haberer, G., Kassa, M.T., Brinton, J., Ramirez-Gonzalez, R.H., Kolodziej, M.C., Delorean, E., Thambugala, D., et al. (2020). Multiple wheat genomes reveal global variation in modern breeding. Nature 588, 277–283.

Wan, H., Li, J., Ma, S., Yang, F., Chai, L., Liu, Z., Wang, Q., Pu, Z., and Yang, W. (2022). Allopolyploidization increases genetic recombination in the ancestral diploid D genome during wheat evolution. Crop J 10, 743–753.

Wang, J., Luo, M.C., Chen, Z., You, F.M., Wei, Y., Zheng, Y., and Dvorak, J. (2013). Aegilops tauschii single nucleotide polymorphisms shed light on the origins of wheat D-genome genetic diversity and pinpoint the geographic origin of hexaploid wheat. New Phytol 198, 925–937.

Wang, M., Li, Z., Zhang, Y., Zhang, Y., Xie, Y., Ye, L., Zhuang, Y., Lin, K., Zhao, F., Guo, J., et al. (2021). An atlas of wheat epigenetic regulatory elements reveals subgenome divergence in the regulation of development and stress responses. Plant Cell 33, 865–881.

Wouters, F.C., Blanchette, B., Gershenzon, J., and Vassao, D.G. (2016). Plant defense and herbivore counter-defense: benzoxazinoids and insect herbivores. Phytochem Rev 15, 1127–1151.

Yang, C., Yang, Z., Zhao, L., Sun, F., and Liu, B. (2018). A newly formed hexaploid wheat exhibits immediate higher tolerance to nitrogen-deficiency than its parental lines. BMC Plant Biol 18, 113.

Yang, C., Zhao, L., Zhang, H., Yang, Z., Wang, H., Wen, S., Zhang, C., Rustgi, S., von Wettstein, D., and Liu, B. (2014). Evolution of physiological responses to salt stress in hexaploid wheat. Proc Natl Acad Sci U S A 111, 11882–11887.

Yonekura-Sakakibara, K., Nakabayashi, R., Sugawara, S., Tohge, T., Ito, T., Koyanagi, M., Kitajima, M., Takayama, H., and Saito, K. (2014). A flavonoid 3-O-glucoside:2"-O-glucosyltransferase responsible for terminal modification of pollen-specific flavonols in Arabidopsis thaliana. Plant J 79, 769–782.

Yu, Z., Duan, X., Luo, L., Dai, S., Ding, Z., and Xia, G. (2020). How plant hormones mediate salt stress responses. Trends Plant Sci 25, 1117–1130.

Yuan, J., Jiao, W., Liu, Y., Ye, W., Wang, X., Liu, B., Song, Q., and Chen, Z.J. (2020). Dynamic and reversible DNA methylation changes induced by genome separation and merger of polyploid wheat. BMC Biol 18, 171.

Yuan, J., Sun, H., Wang, Y., Li, L., Chen, S., Jiao, W., Jia, G., Wang, L., Mao, J., Ni, Z., et al. (2022). Open chromatin interaction maps reveal functional regulatory elements and chromatin architecture variations during wheat evolution. Genome Biol 23, 34.

Zhang, F., Wu, J., Sade, N., Wu, S., Egbaria, A., Fernie, A.R., Yan, J., Qin, F., Chen, W., Brotman, Y., et al. (2021a). Genomic basis underlying the metabolome-mediated drought adaptation of maize. Genome Biol 22, 260.

Zhang, H., Zhu, B., Qi, B., Gou, X., Dong, Y., Xu, C., Zhang, B., Huang, W., Liu, C., Wang, X., et al. (2014). Evolution of the BBAA component of bread wheat during its history at the allohexaploid level. Plant Cell 26, 2761–2776.

Zhang, Y., Liu, T., Meyer, C.A., Eeckhoute, J., Johnson, D.S., Bernstein, B.E., Nusbaum, C., Myers, R.M., Brown, M., Li, W., et al. (2008). Model-based Analysis of ChIP-Seq (MACS). Genome Biol 9, R137.

Zhang, Z., Tan, M., Zhang, Y., Jia, Y., Zhu, S., Wang, J., Zhao, J., Liao, Y., and Xiang, Z. (2021b). Integrative analyses of targeted metabolome and transcriptome of Isatidis Radix autotetraploids highlighted key polyploidization-responsive regulators. BMC Genomics 22, 670.

Zhao, J., and Dixon, R.A. (2010). The ’ins’ and ’outs’ of flavonoid transport. Trends Plant Sci 15, 72-80.

Zhao, L., Wang, S., Cao, Z., Ouyang, W., Zhang, Q., Xie, L., Zheng, R., Guo, M., Ma, M., Hu, Z., et al. (2019). Chromatin loops associated with active genes and heterochromatin shape rice genome architecture for transcriptional regulation. Nat Commun 10, 3640.

Zhao, L., Yang, Y., Chen, J., Lin, X., Zhang, H., Wang, H., Wang, H., Bie, X., Jiang, J., Feng, X., et al. (2023). Dynamic chromatin regulatory programs during embryogenesis of hexaploid wheat. Genome Biol 24, 7.

Zhao, L., Xie, L., Zhang, Q., Ouyang, W., Deng, L., Guan, P., Ma, M., Li, Y., Zhang, Y., Xiao, Q., et al. (2020). Integrative analysis of reference epigenomes in 20 rice varieties. Nat Commun 11, 2658.

Zhou, Y., Zhao, X., Li, Y., Xu, J., Bi, A., Kang, L., Xu, D., Chen, H., Wang, Y., Wang, Y.-g., et al. (2020). Triticum population sequencing provides insights into wheat adaptation. Nat Genet 52, 1412–1422.

