## Supplementary Fig. S1-S18 for "Integration of the DD-genome reshapes gene transcription, chromatin architecture and metabolome of allohexaploid wheat leading to enhanced adaptability"

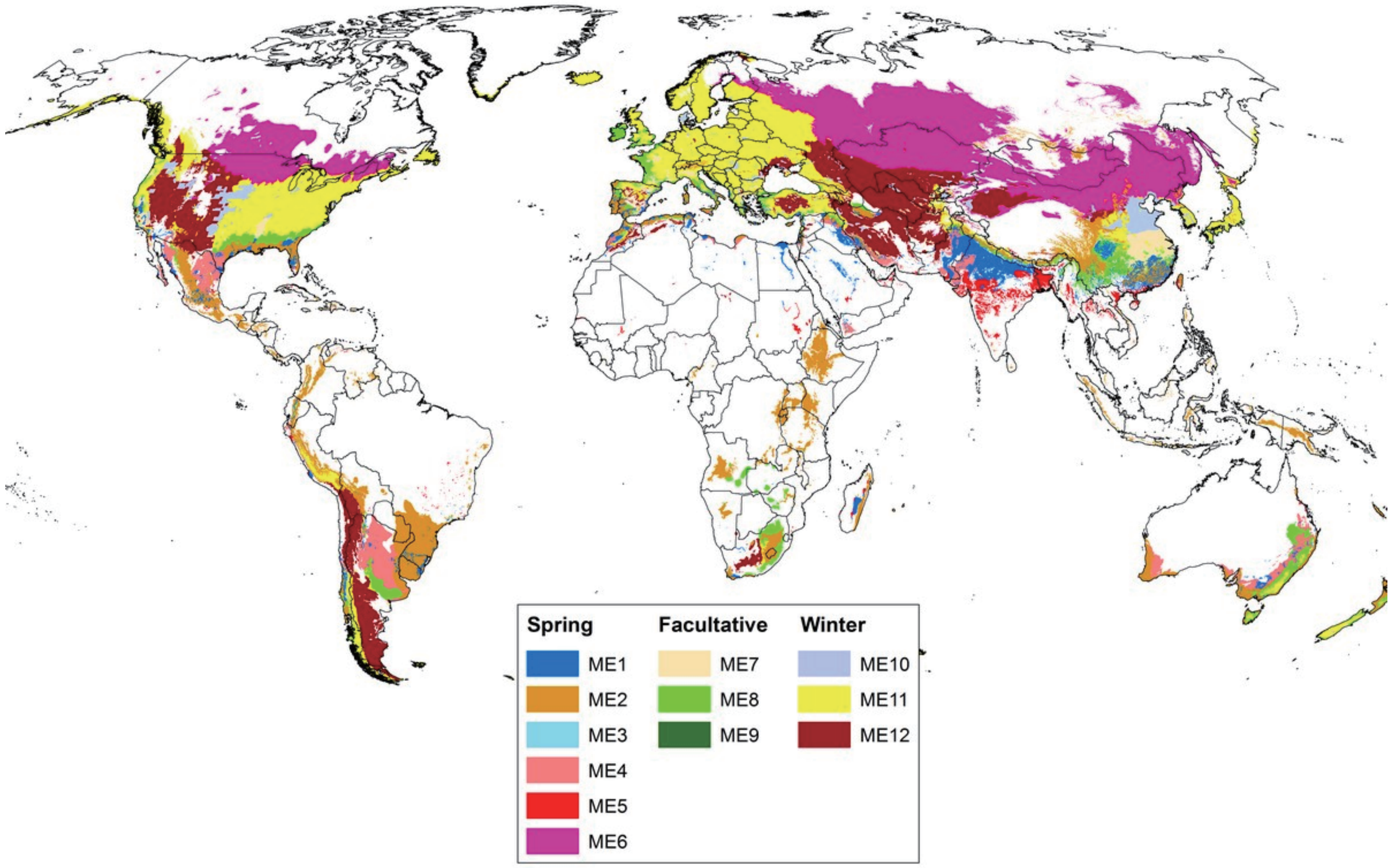


**Supplementary Figure S1. Global Distribution of Wheat Megaenvironments.** Different colors represent different mega-environments (ME) that showed different biotic and abiotic stresses and cropping systems. ME1: low rainfall, irrigated, temperature (T) for coolest quarter 11°C > T ≥ 3°C. Major constraints: leaf rust, stem rust, stripe rust, lodging. ME2: high rainfall, temperature for coolest quarter 16°C > T ≥ 3°C. Major constraints: barley yellow dwarf virus, stripe rust, head scab, septoria, tan spot. ME3: high rainfall, temperature for coolest quarter 16°C > T ≥ 3°C, soil pH < 5.2. Major constraints: barley yellow dwarf virus, stripe rust, head scab, septoria, tan spot, acid soil (Aluminum, Manganese toxicity). ME4: low rainfall, temperature for coolest quarter 16°C > T ≥ 3°C. Major constraints: drought, rust, common bunt, septoria, fusarium, tan spot, root diseases. ME5: tropical high rainfall, and/or irrigated, temperature for coolest quarter 16°C > T > 11°C. Major constraints: heat, spot blotch, leaf & stem rust. ME6: high latitude (>45° N or S), temperature for coolest quarter T < -13°C and for warmest quarter T = 9°C. Major constraints: rust, root diseases, tan spot. ME7: irrigated, temperature for coolest quarter 3°C > T ≥ -2°C. Major constraints: cold, stripe rust, mildew. ME8: high rainfall/irrigated, temperature for coolest quarter 6°C > T ≥ 1°C. Major constraints: cold, stripe rust, mildew, septoria, root rots. ME9: low rainfall, temperature for coolest quarter 3°C > T ≥ -2°C. Major constraints: cold, drought, stripe rust, root rots. ME10: irrigated, temperature for coolest quarter -2°C > T ≥ -13°C. Major constraints: winter kill, rust, mildew. ME11: high rainfall/irrigated, temperature for coolest quarter 1°C > T ≥ -13°C. Major constraints: winter kill, rust, septoria, mildew. ME12: low rainfall, temperature for coolest quarter 1°C > T ≥ -13°C. Major constraints: winter kill, drought, stripe rust, bunts, root rots. The data was download from website http://wheatatlas.org/me-global-distribution.

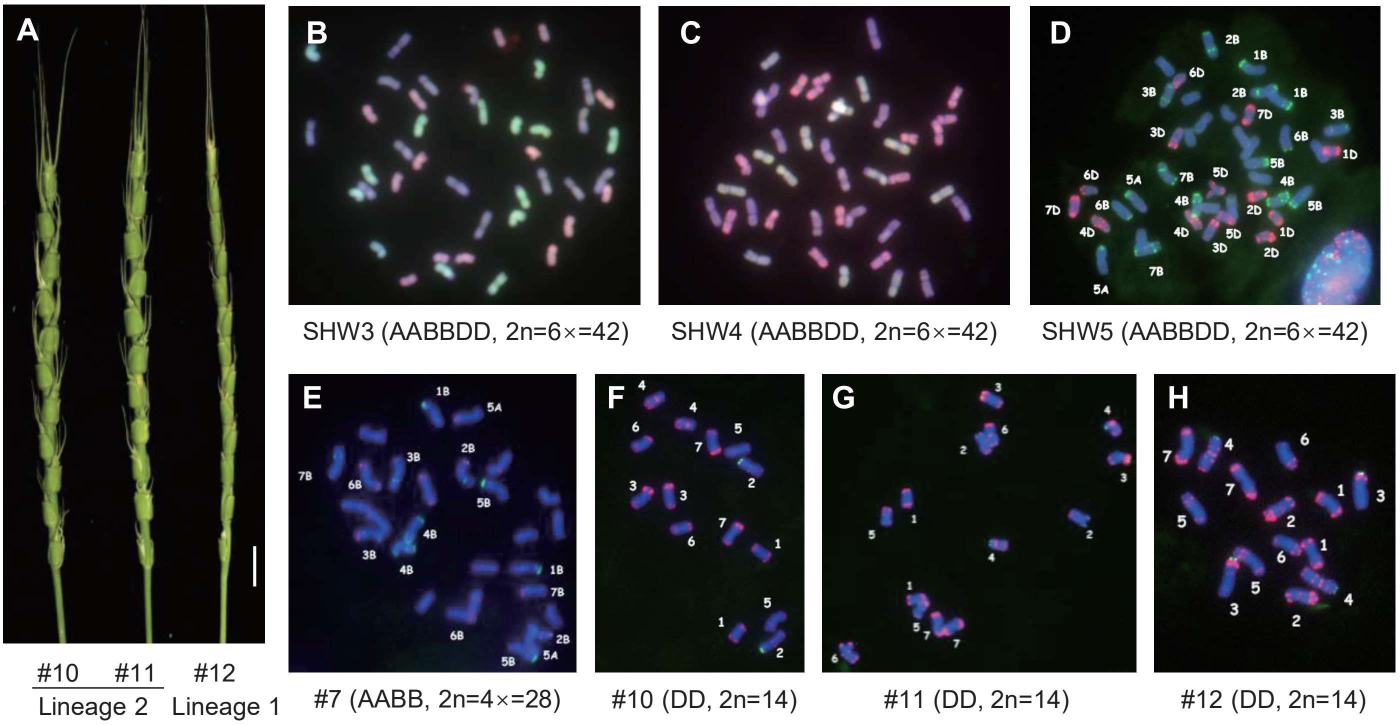


**Supplementary Figure S2. Morphological characteristics of the diploid *Ae. tauschii* and cytology karyotyping analysis of materials used in this study.**

**(A)** Morphological characteristics of spikes of the diploid parents used in this study. Bar = 1 cm. **(B, C)** Genomic in situ hybridization of mitotic cells in the root tip from allohexaploid SHW3 (**B**), SHW4 (**C**) colored by the subgenome A (green), B (blue), D (pink). **(D-H)** Fluorescent in situ hybridization using mitotic cells of the root tip from allohexaploid SHW5 (**D**), tetraploid parent *T. turgidum* (**E**) and diploid parents *Ae. tauschii* (**F-H**) using pAS1-1 (red) and 119.2 (green) as probes.

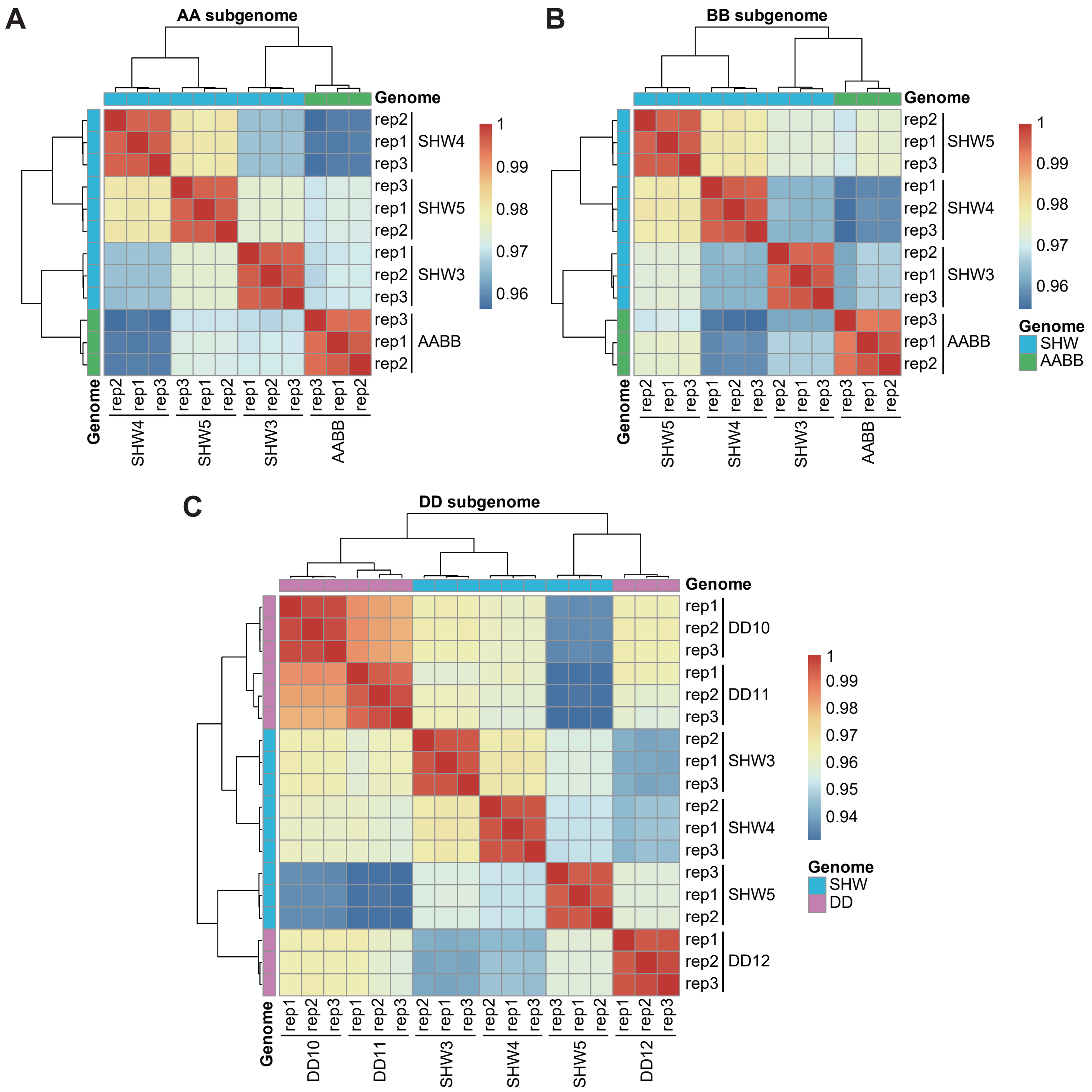


**Supplementary Figure S3. Reproducibility analysis of transcriptome data in this study.**
**(A)** Correlation coefficients of A subgenome transcriptome data from materials containing A genome.
**(B)** Correlation coefficients of B subgenome transcriptome data from materials containing B genome.
**(C)** Correlation coefficients of D subgenome transcriptome data from materials containing D genome.

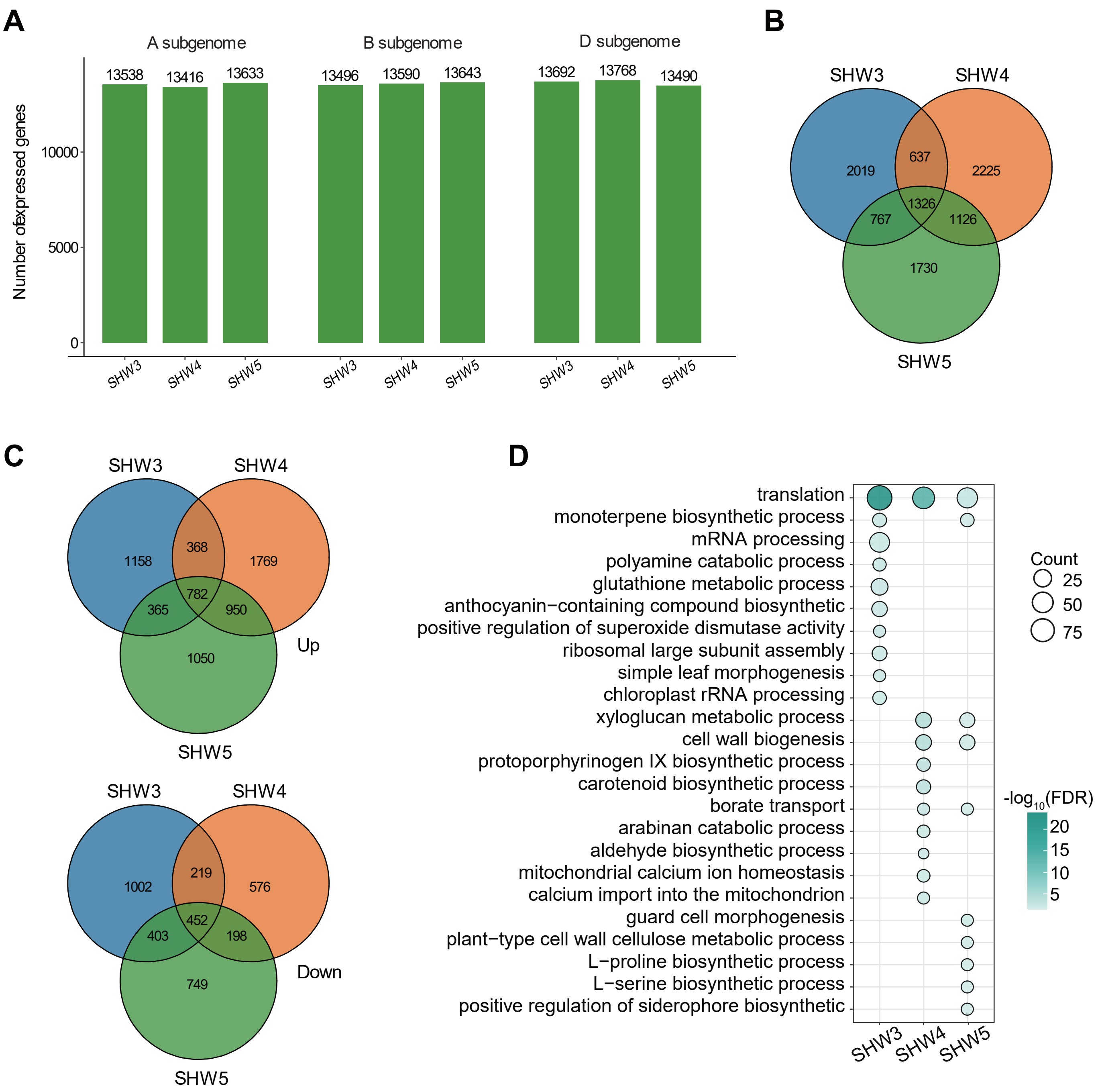


**Supplementary Figure S4. Gene expression analysis.**
**(A)** The total number of expressed genes in SHW3, SHW4 and SHW5. Genes were considered to be expressed only when their expression level tpm > 0.5. **(B)** Venn diagrams showing the overlap of DEGs in SHW3, SHW4 and SHW5. **(C)** Venn diagrams showing the overlap of activated genes (top) and suppressed genes (bottom) in SHW3, SHW4 and SHW5. **(D)** Top ten GO terms enriched in down-regulated DEGs. Dot size represents the gene number enriched in GO terms and color gradation represents the significance of enrichment as $-\log_{10}$ (FDR).

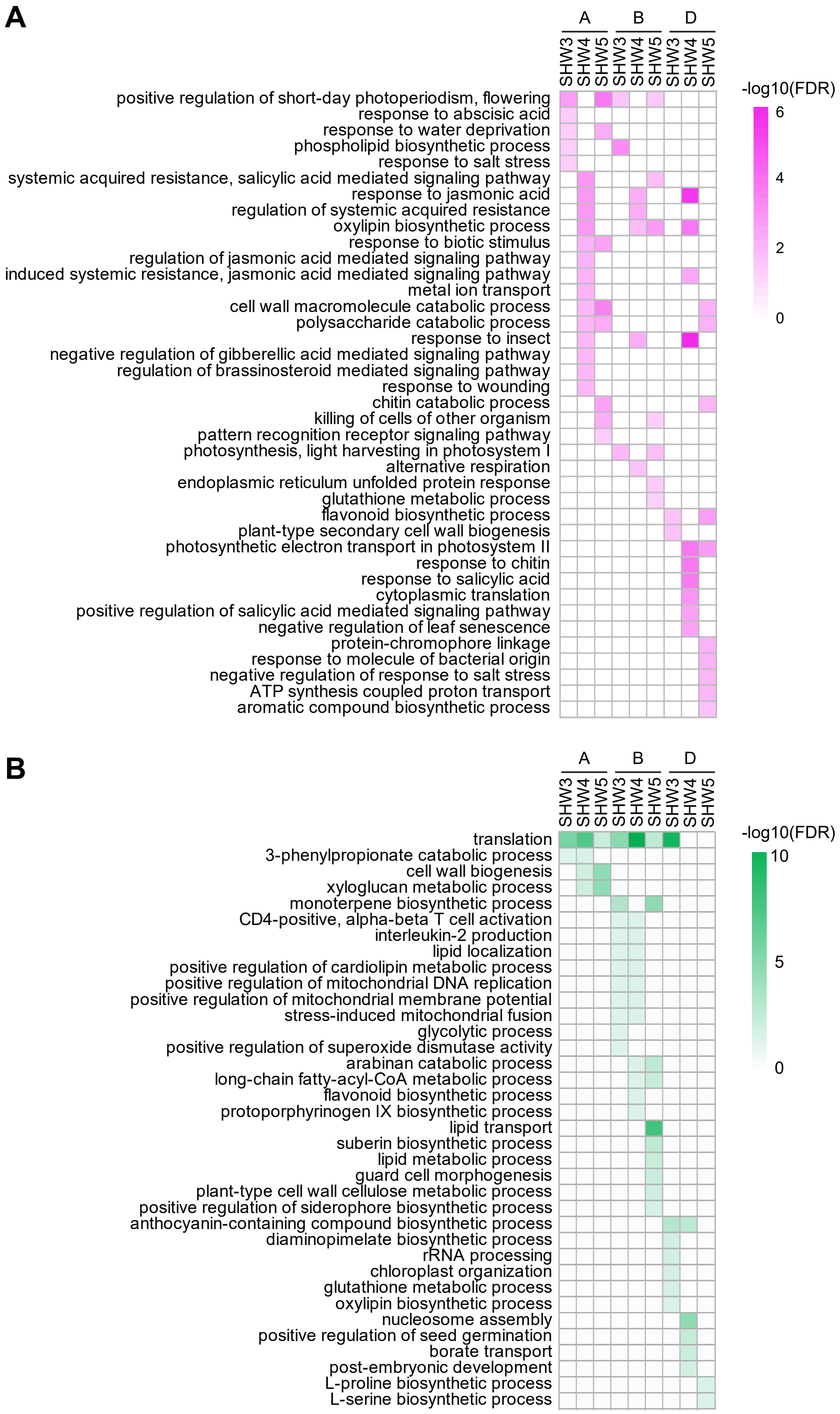


**Supplementary Figure S5. GO enrichment analysis for DEGs in different subgenomes.**
**(A)** Enriched biological process GO terms for up-regulated DEGs in SHW3, SHW4 and SHW5. **(B)** Enriched biological process GO terms for down-regulated DEGs in SHW3, SHW4 and SHW5. The gradation of color represents the significance of GO terms as $-\log_{10}$ (FDR).

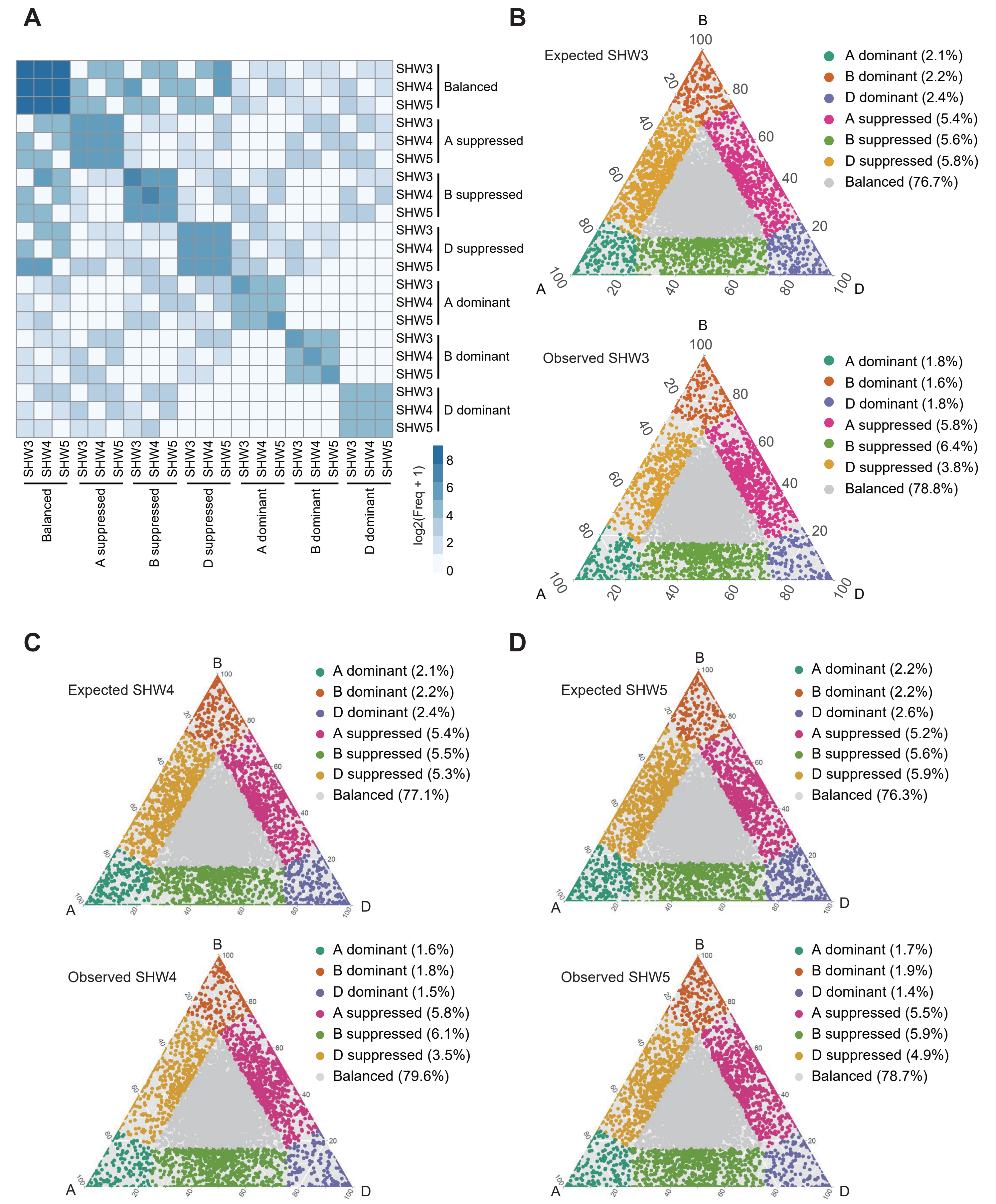


**Supplementary Figure S6. Features of the seven categories of gene triads.**
**(A)** Heatmaps showing relations of different categories of SHW3, SHW4 and SHW5. **(B-D)** Ternary plots showing the expression bias of homoeolog triads in SHW3 (**B**), SHW4 (**C**) and SHW5 (**D**) before (expected, top) and after (observed, bottom) polyploidization. Each point indicates a gene triad with relative contribution of A, B and D homoeologs to an overall triad expression. Each of the seven different categories were shown by a particular color. The proportion of seven categories were shown in brackets.

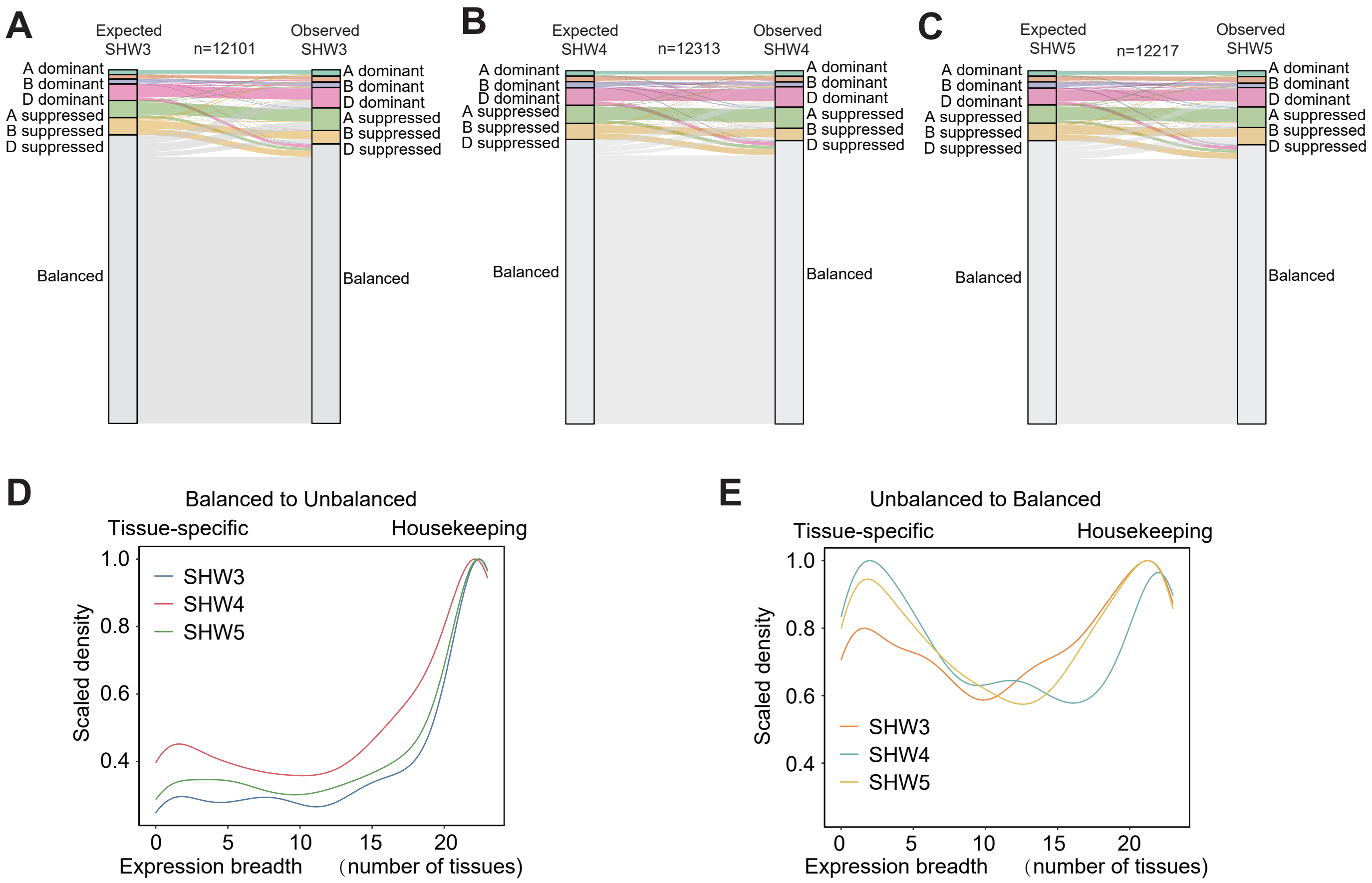


**Supplementary Figure S7. Asymmetric expression changes of subgenome homoeologs after polyploidization.**

**(A-C)** Alluvial plots showing the reshaping of expression asymmetry of homoeologs in SHW3 (**A**), SHW4 (**B**) and SHW5 (**C**) during polyploidization. Only triads expressed both before and after polyploidization were shown. **(D, E)** Expression breadth analysis of genes in triads showed balanced-to-unbalanced (**D**) or unbalanced-to-balanced (**E**) transition during polyploidizations. The more tissues in which a gene is expressed, the higher chance the gene is a housekeeping gene; while a gene is expressed in a few tissue types, it is a tissue-specific gene. Different color lines showing different SHWs.

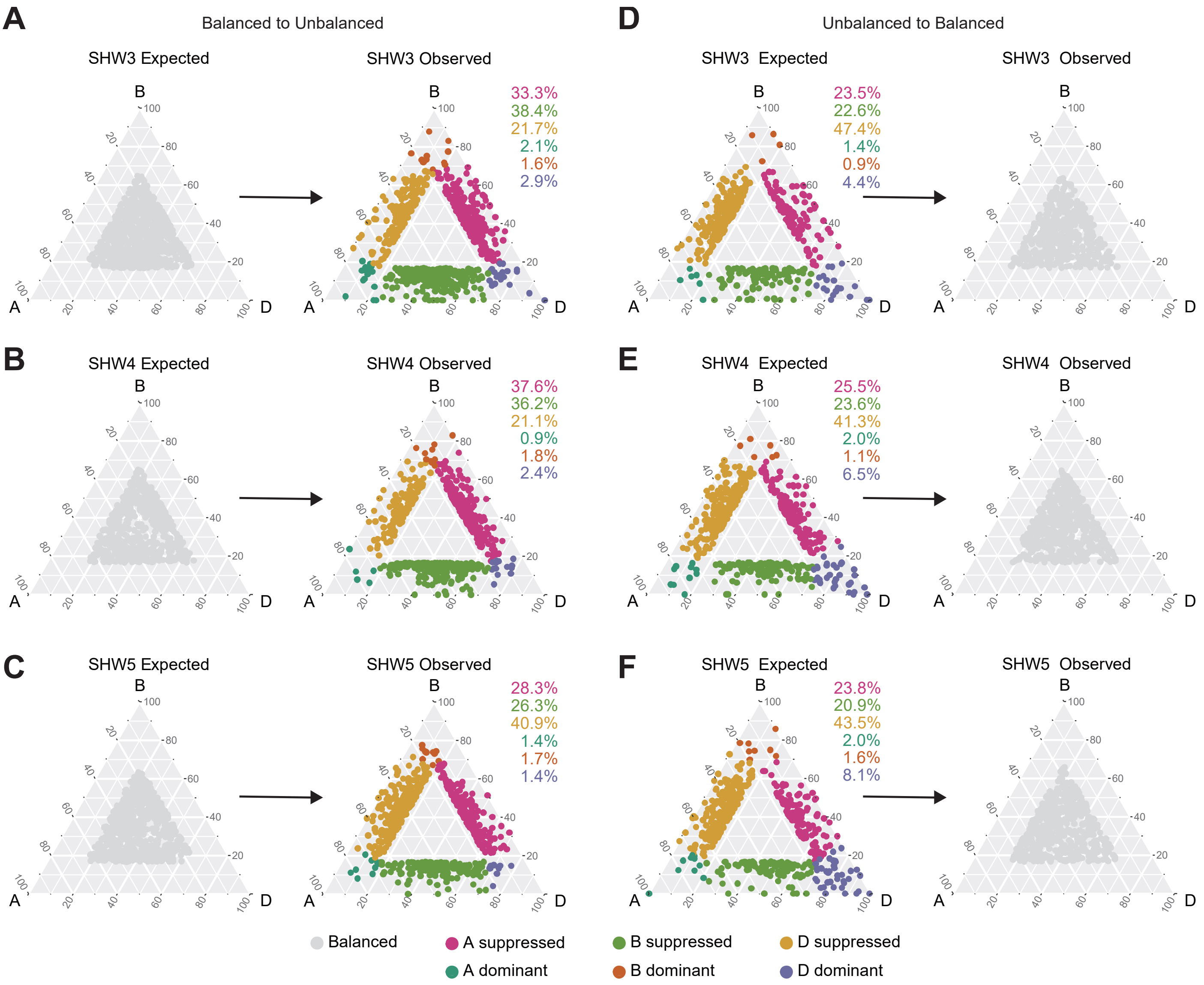


**Supplementary Figure S8. Ternary plots showing triads that altered its balancing state during polyploidizations.**

**(A-C)** Points in ternary plots represent triads that showed expression bias state from balanced to unbalanced categories after polyploidizations in SHW3 (**A**), SHW4 (**B**) and SHW5 (**C**). **(D-F)** Points in ternary plots indicate triads which showed expression bias state from unbalanced to balanced in SHW3 (**D**), SHW4 (**E**) and SHW5 (**F**). Points belong to different triad categories are shown with different colors. The proportion of unbalanced triad is giving in same color with corresponding circles.

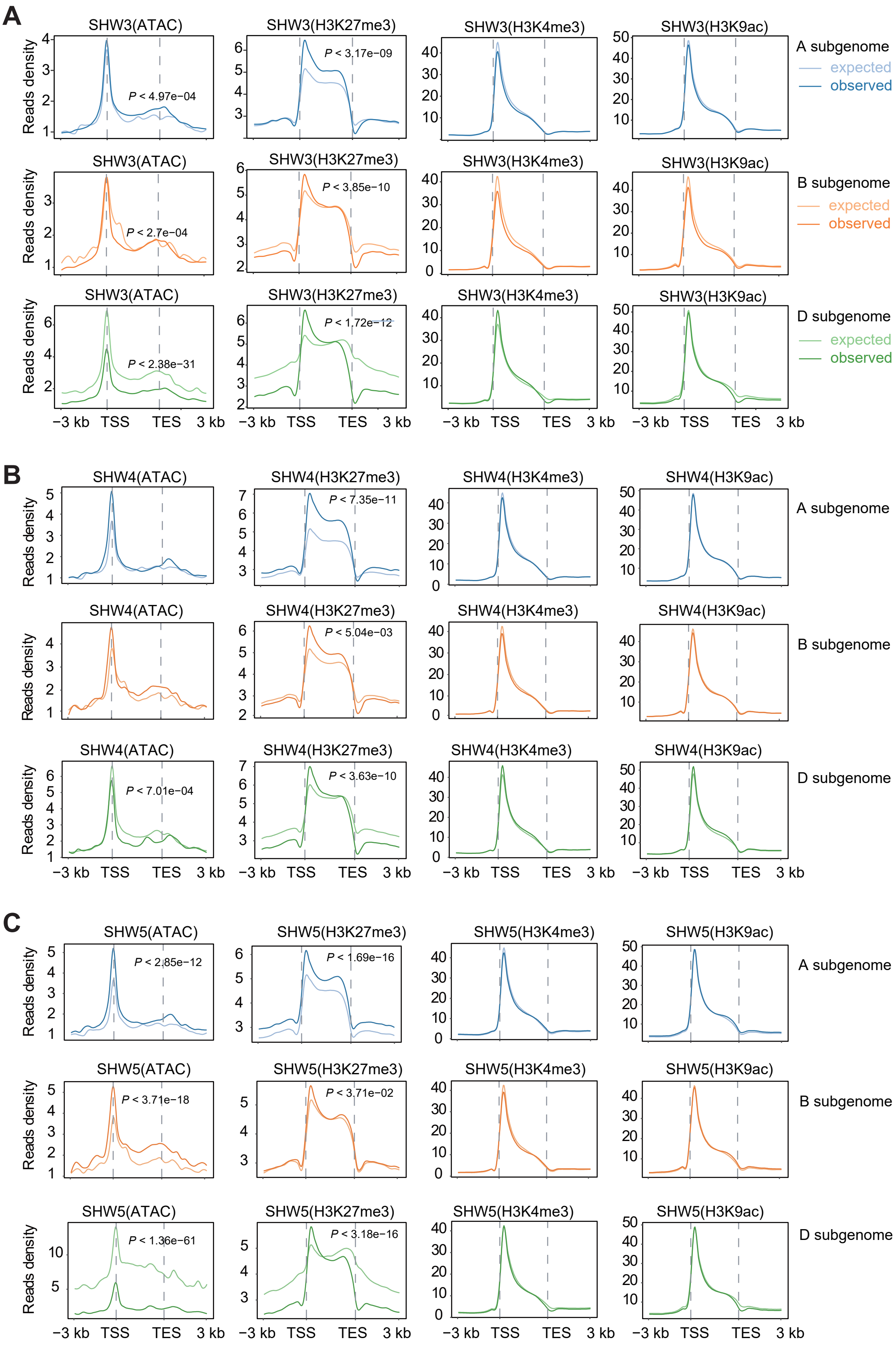


**Supplementary Figure S9. Comparing epigenomic changes in SHWs before and after polyploidization.**

**(A-C)** Chromatin accessibility (ATAC-seq) and histone modification levels of H3K27me3, H3K4me3, H3K9ac in A (top), B (middle) and D (bottom) subgenomes of SHW3 (**A**), SHW4 (**B**) and SHW5 (**C**). Light lines represent reads density before polyploidization (expected); dark lines indicate reads density after polyploidization (observed).

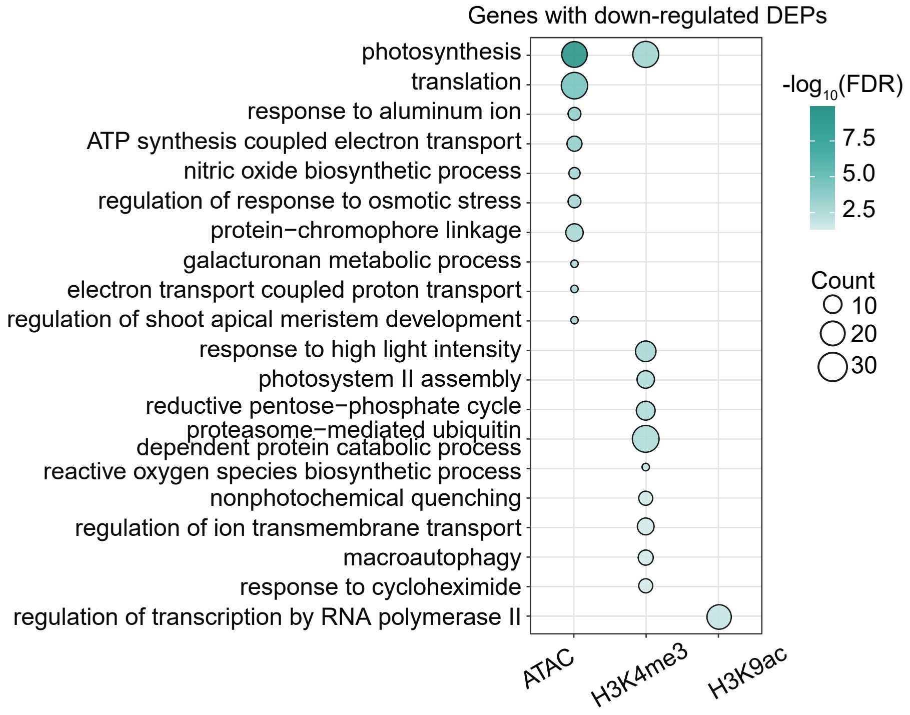


**Supplementary Figure S10. GO enrichment analysis of target genes with down-regulated DEPs.** Color gradation indicates the significance as $-\log_{10}$(FDR). Dot size represents gene number enriched in that term.

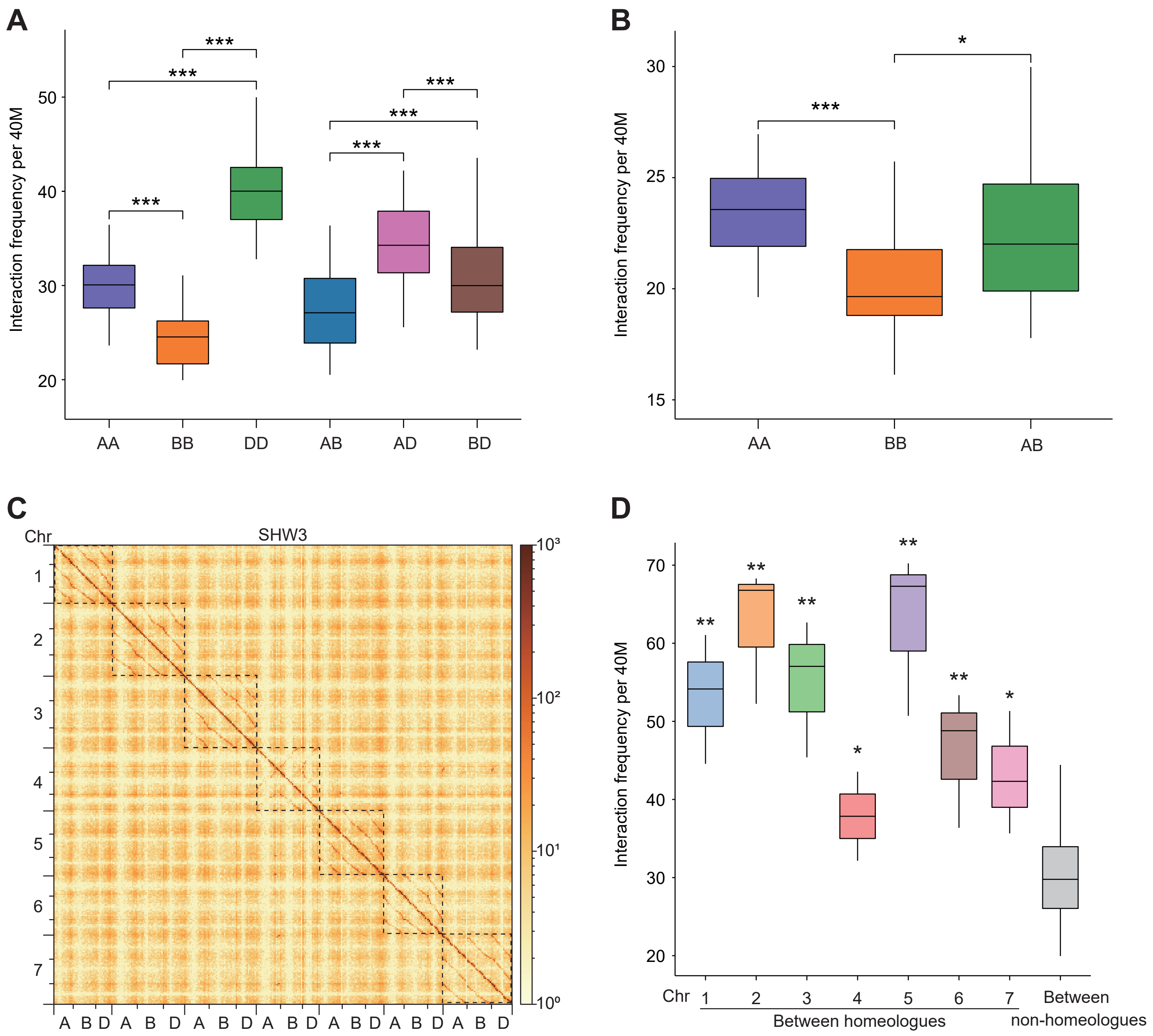


**Supplementary Figure S11. Analysis of H3K4me3-associated 3D chromatin interactions in SHW3 and its tetraploid parent.**

**(A, B)** Boxplots representing interaction frequency within or between subgenomes in SHW3 (**A**) and its tetraploid parent (**B**). **(C)** Heatmap showing chromatin interactions between homeologues associated with H3K4me3 modification in SHW3. **(D)** Boxplots showing interaction frequency between homoeologous chromosomes comparing to non-homoeologous chromosomes of SHW3. **P* < 0.05, ***P* < 0.01, ****P* < 0.001.

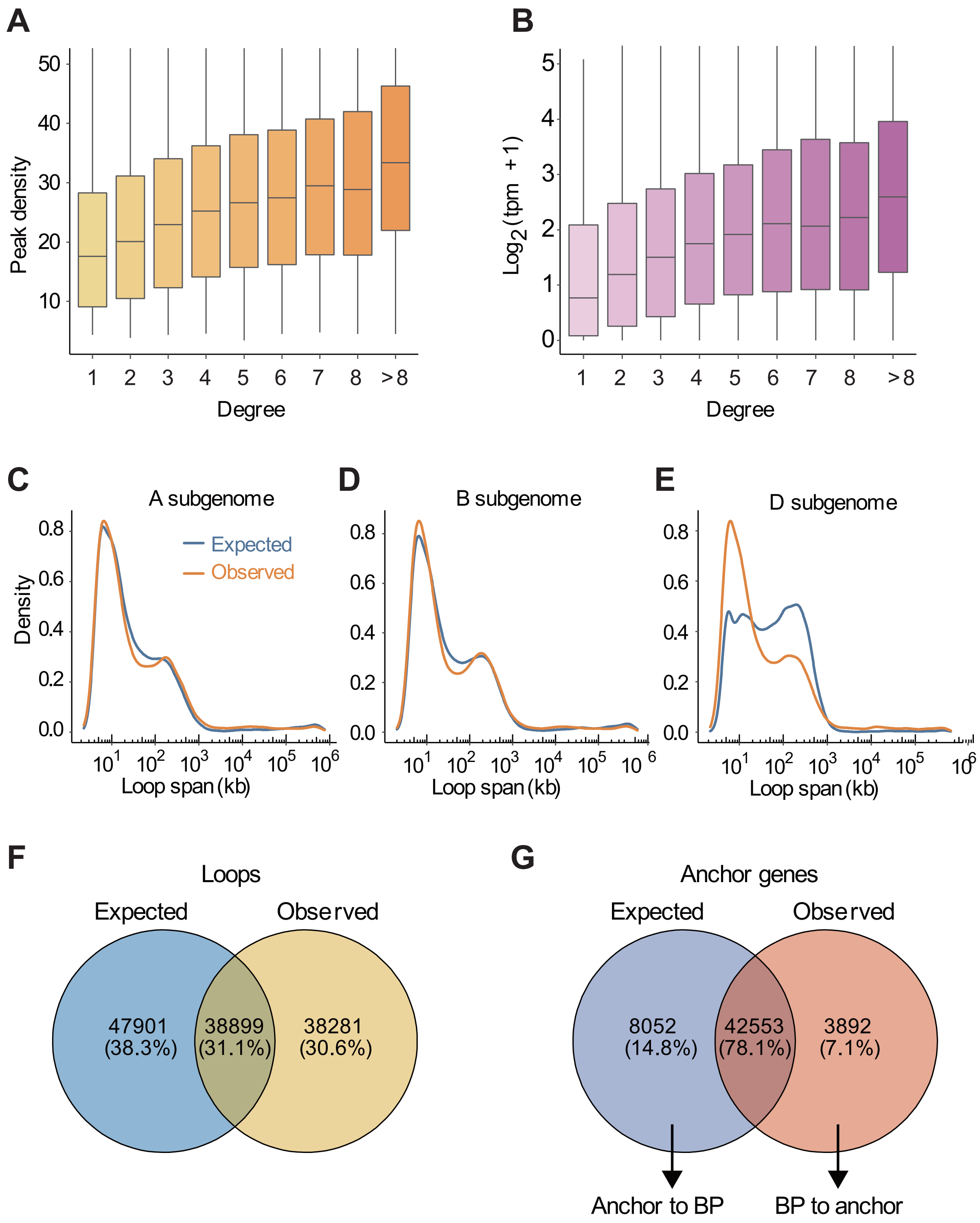


**Supplementary Figure S12. Characterization of H3K4me3-associated chromatin interactions in SHW3 and its parents.**

**(A, B)** Boxplots showing the positive relationship of H3K4me3 peak intensity (**A**) and expression level (**B**) with anchor gene degree in SHW3. Expression levels showing as $\log_2$(tpm + 1). **(C-E)** Genomic span distribution of intrachromosomal interaction loops in A (**C**), B (**D**) and D (**E**) subgenomes of SHW3 before (expected) and after (observed) polyploidization. **(F, G)** Venn diagram showing overlaps of interaction loops (**F**) and anchor genes (**G**) in expected and observed dataset.

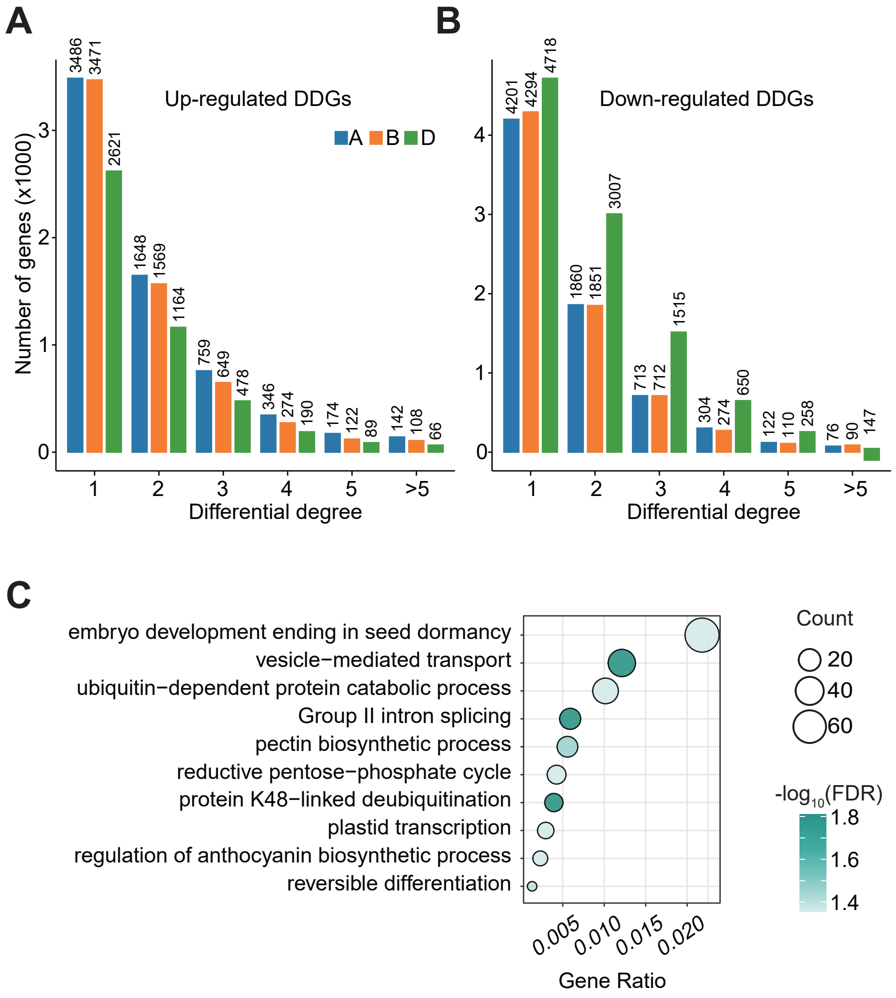


**Supplementary Figure S13. Analysis of genes showed altered interaction degree during wheat polyploidization.**

**(A, B)** Statistics of up-regulated differential degree genes (DDGs) **(A)** and down-regulated DDGs **(B)** in different subgenomes during wheat polyploidization. **(C)** Bubble diagram showing enriched GO terms of down-regulated DDGs in SHW3. The dot size represents the number of genes in the corresponding GO term, and the gradation of color indicates the significance of terms as $-\log_{10}$ (FDR).

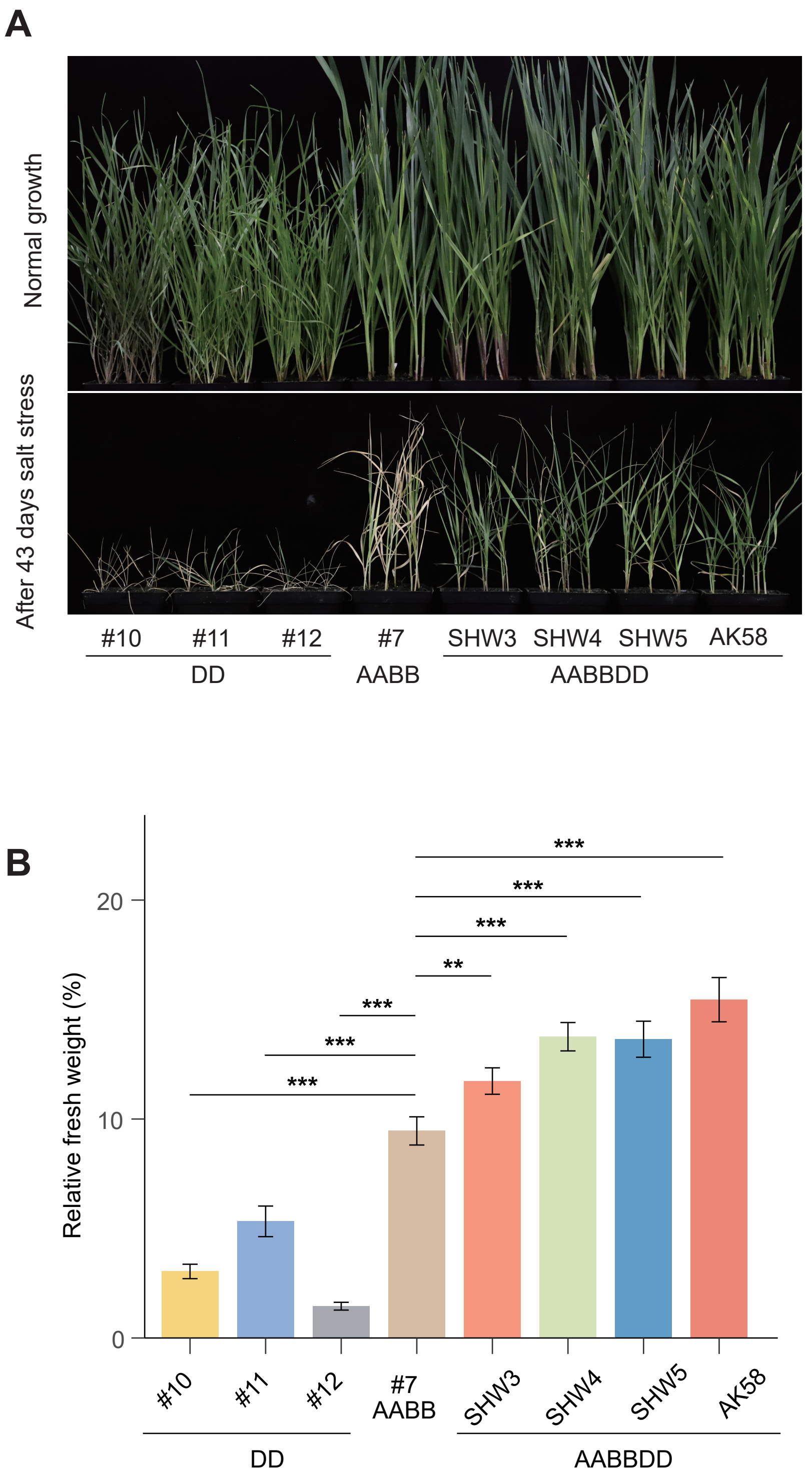


**Supplementary Figure S14. Phenotypes of AK58, SHWs, and tetraploid and diploid parents (AABB and DD) under salt stress.**

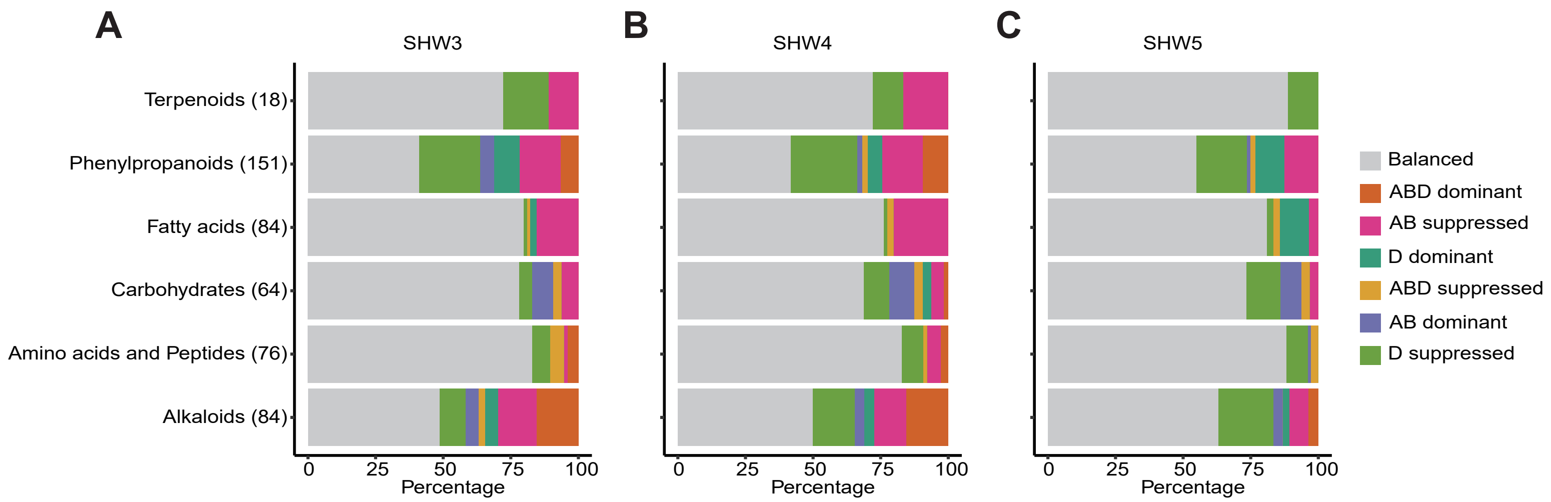


**Supplementary Figure S15. Comparation of metabolome in SHWs and its parents.**
**(A-C)** The percentage of seven categories of content balance between SHWs and their tetraploid, diploid parents of SHW3 (**A**), SHW4 (**B**), SHW5 (**C**) in metabolite classes with annotated structure.

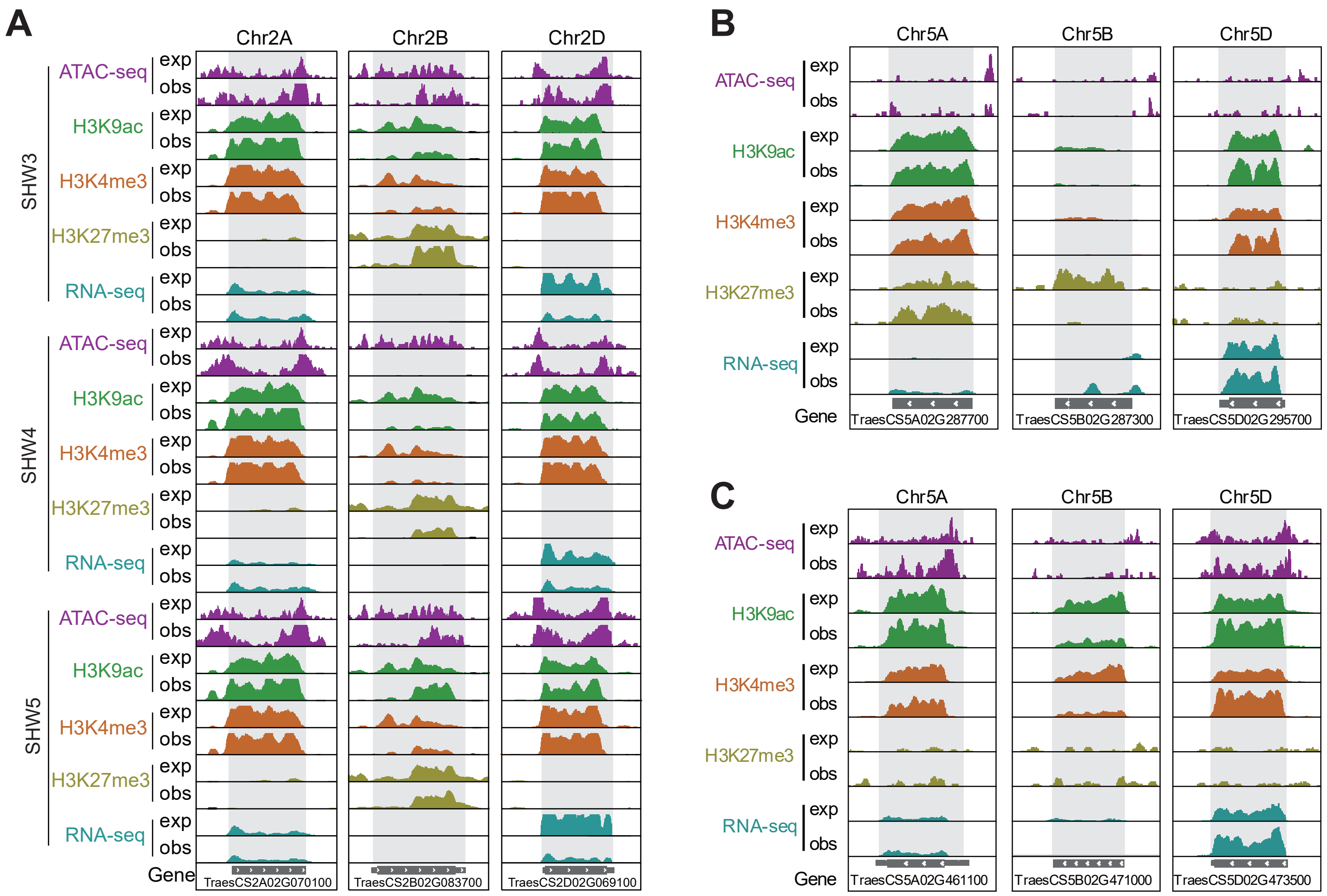


**Supplementary Figure S16. Epigenomic features of flavonoid pathway genes during DD genome integration-mediated polyploidization.**

**(A)** Epigenomic features of *TraesCS2D01G069100* homoeologs in SHW3, SHW4 and SHW5 before (exp) or after (obs) DD-integration events. **(B, C)** Epigenomic features of subgenome homoeolog genes of *TraesCS5D02G295700* (**B**) and *TraesCS5D02G473500* (**C**) in SHW3 during polyploidization. The highlighted regions in grey representing the gene body.

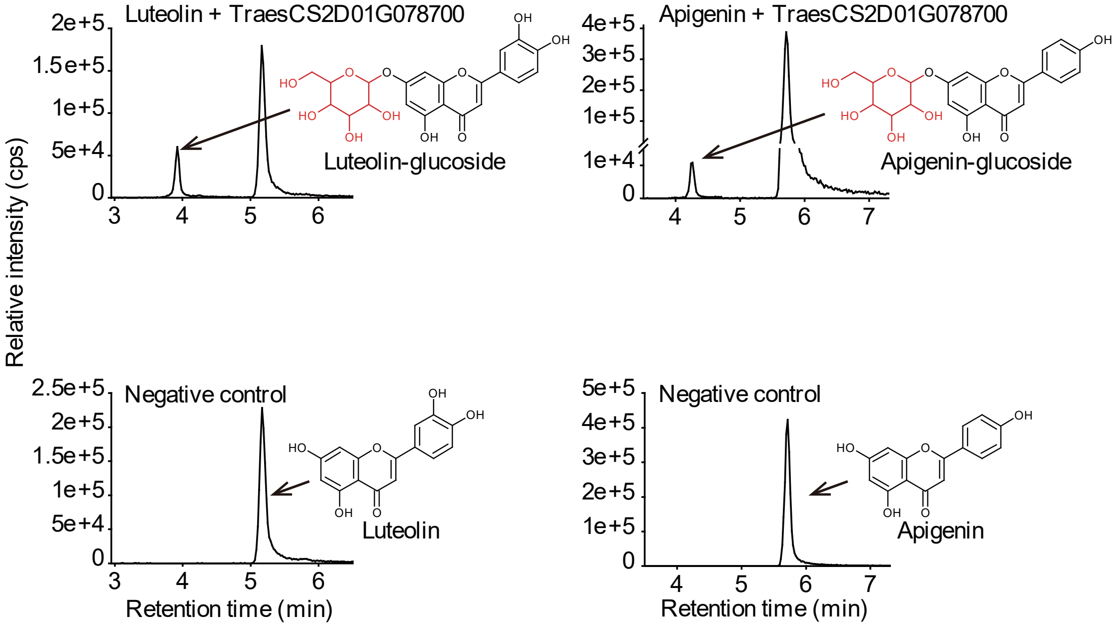


**Supplementary Figure S17. The O-glucosyltransferase activities of TraesCS2D01G078700 from D genome. Newly transferred glycosyl is shown in red color.**

A

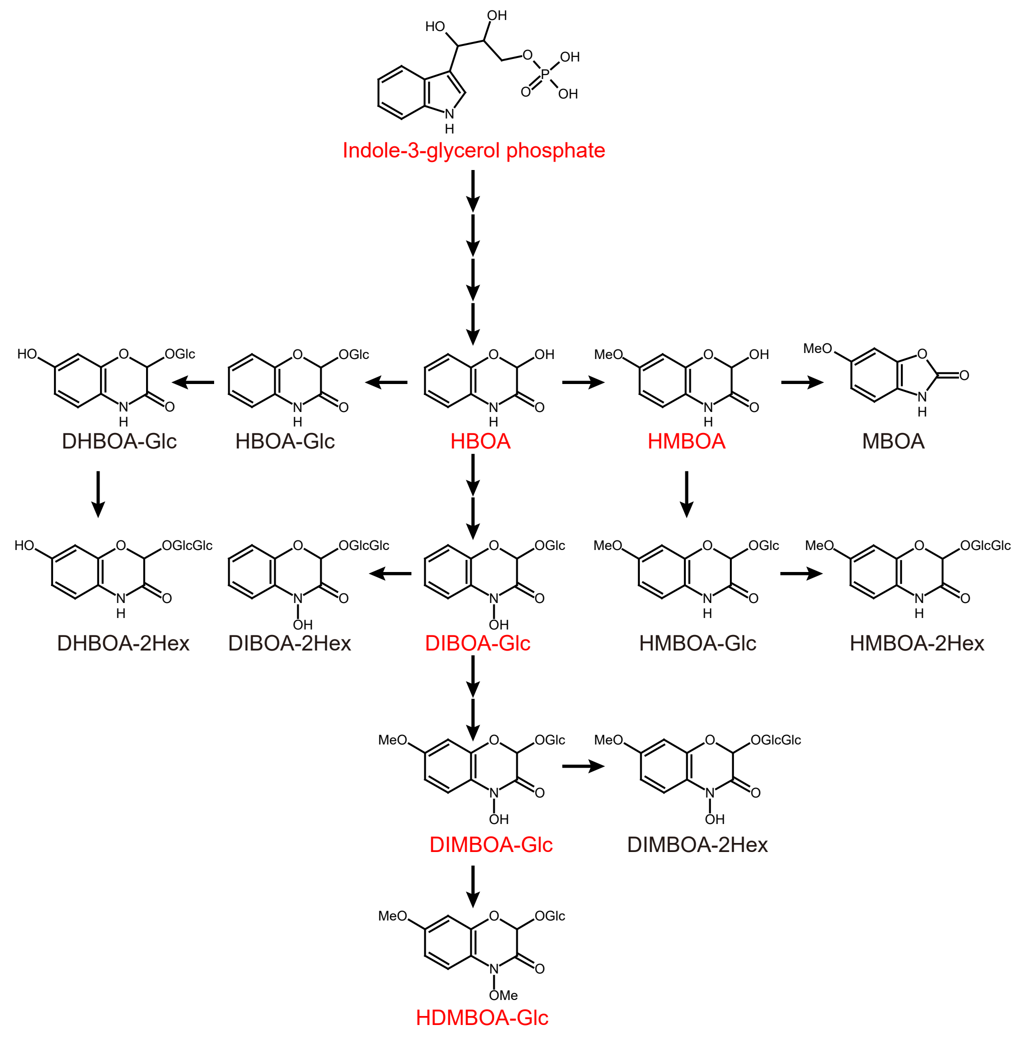


B

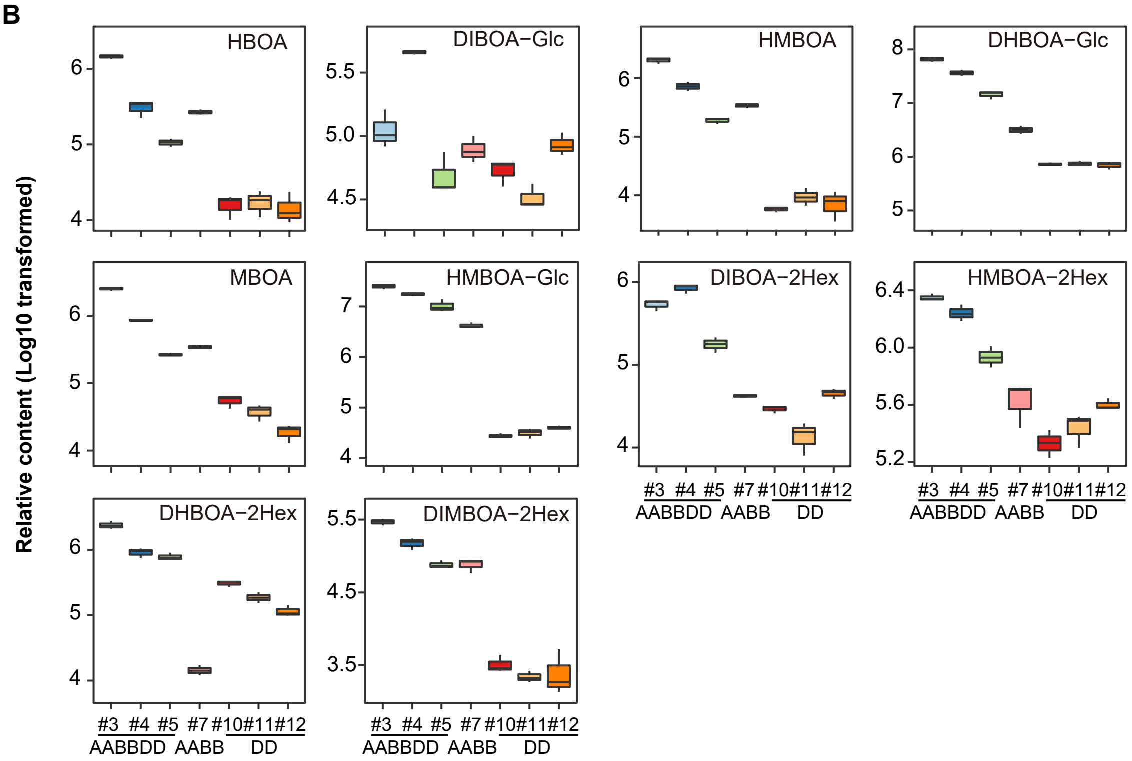


**Supplemental Figure S18. Metabolites in benzoxazinoid pathway detected in this study.**
**(A)** Supplementary of Figure 6F. The putative biosynthesis pathway of benzoxazinoid derivatives detected in this study. Red color indicates core metabolites shown in Figure 6F. **(B)** Relative contents of benzoxazinoid derivatives in hexaploid SHWs and its tetraploid and dipoid parents. Contents are given as $log_{10}$ transformed.
